# VGLL1 contributes to both the transcriptome and epigenome of the developing trophoblast compartment

**DOI:** 10.1101/2025.04.25.650693

**Authors:** Ruben I. Calderon, Nirvay Sah, Molly Huang, Sampada Kallol, Ryan H. Kittle, Walee B. Shaik, Ahmed Abdelbaki, Jennifer N. Chousal, Robert Morey, Tony Bui, Alejandra Mitre, Norah M.E. Fogarty, Claudia Gerri, Zoe Manalo, Claire Zheng, Peter De Hoff, Pratik Home, Kathy K. Niakan, Heidi Cook-Andersen, Kathleen M. Fisch, Soumen Paul, Francesca Soncin

## Abstract

The trophectoderm (TE), the first lineage specified during mammalian development, initiates implantation and gives rise to placental trophoblasts. While animal models have elucidated key conserved signaling pathways involved in early TE specification, including BMP, WNT, and HIPPO, species-specific differences during early development emphasize the need for human-specific models. We previously identified VGLL1, a coactivator of TEAD transcription factors, as a human-specific placental marker. In this study, we employed a pluripotent stem cell (PSC)-based model of TE induction by BMP4 to investigate chromatin remodeling and transcriptional dynamics during TE formation. BMP4-induced chromatin accessibility changes promoted a trophoblast gene expression program, while mesoderm lineage markers were only transiently expressed upon canonical WNT activation. We found that VGLL1 was expressed downstream of key TE transcription factors (GATA2/3, TFAP2A/C) but was essential for establishment of full trophoblast identity by up-regulating EGFR and reinforcing GATA3 expression through positive feedback. Notably, VGLL1 enhanced canonical WNT signaling via direct regulation of WNT receptors and effectors. We also identified KDM6B, a histone demethylase that removes H3K27me3 repressive marks, as a direct VGLL1 target. KDM6B facilitated activation of bivalent promoters associated with TE markers, linking epigenetic regulation to lineage identity. Our findings establish a mechanistic framework positioning VGLL1 as a central regulator that integrates HIPPO, BMP, and WNT signaling pathways to drive establishment of human TE.

**Statement of Significance:** Early development of the human placenta is essential for pregnancy success, yet the mechanisms that guide placental lineage specification remain poorly defined. Using human stem cells, we show how signaling pathways and chromatin remodeling programs work together to direct formation of the trophectoderm, the earliest placental cell type. We identify VGLL1 as a key regulator linking multiple signaling networks to gene expression and epigenetic control. Our findings reveal a species-specific mechanism of placental initiation with broad implications for understanding reproductive disorders, pregnancy loss, and advancing stem cell–based models to study and potentially treat human placental disease.

## Introduction

The trophectoderm (TE) is the first lineage specified during mammalian development and is important for embryo cavitation, implantation, and pregnancy initiation (1). After implantation, the TE gives rise to the trophoblast compartment of the placenta, contributing to the formation of the maternal/fetal interface (2). Early pregnancy loss and several placental-associated disorders are linked to abnormalities in the establishment and differentiation of the TE compartment during early development (2). However, ethical and technical considerations have limited our understanding of the mechanisms of TE specification and trophoblast development in humans. Studies in animal models have determined that tight spatio-temporal regulation of highly conserved signaling pathways, including BMP, WNT, and HIPPO, are required to establish the TE in the pre-implantation embryos and for early placental development (3–8). However, increasing evidence points to species-specific differences in the composition of regulatory networks that contribute to the development of the extra-embryonic ectoderm compartment (3, 9–11). Therefore, species-specific models are better suited to investigate TE specification and early placental development (3, 9, 12).

The HIPPO signaling pathway is a conserved cascade that initiates the specification of the TE lineage following activation of GATA3 and CDX2 expression by the TEAD4/YAP1 complex (10, 13, 14). In early mouse embryos, differential HIPPO pathway activation between the inner and outer cells leads to the expression of CDX2, which initiates a TE expression program, including induction of GATA3 and ELF5 (13, 15, 16). EOMES is expressed down-stream of CDX2 and, while not required for TE specification, it is fundamental for the maintenance of the TE transcriptional program and maintenance of mouse trophoblast stem cells (TSCs) (17, 18). In humans, four transcription factors, GATA2, GATA3, TFAP2A, and TFAP2C, form a transcriptional network required for TE initiation (TEtra factors) downstream of the HIPPO signaling pathway (19). This network drives the establishment of an extra-embryonic ectoderm signature, which includes mature markers such as VGLL1, TP63, and EGFR, but not EOMES, which is dispensable for human TE and trophoblast development (9, 20). In particular, we have previously identified VGLL1 as a human-specific marker expressed in the trophoblast progenitor compartments of the placenta (9). VGLL1 is a conserved binding partner of the TEAD family of transcription factors and competes with YAP1 to form a transcriptional complex with TEAD4, therefore potentially modulating HIPPO signaling in human placenta (21).

Canonical WNT signaling is a well-characterized developmental pathway mediated by nuclear localization of β-catenin, where it binds to TCF/LEF complexes and drives expression of WNT-specific target genes (22). This pathway is required for maintenance and expansion of various human epithelial stem cells, including intestinal and skin stem cells (23, 24). In human pre-implantation embryos, the TE compartment is enriched in WNT components, including ligands and receptors (25–29). Moreover, compared to mouse, both human and non-human primate embryos show higher canonical WNT-related gene expression in the TE (30). WNT genes are also highly expressed in the trophoblast compartment of the placenta. *In vitro*, both human TSCs and trophoblast organoids require activation of canonical WNT signaling for proliferation (31–33) while its transient inhibition is required to initiate differentiation of extravillous trophoblasts (EVT) (32, 34).

Bone Morphogenetic Protein (BMP) signaling has well described roles in the embryonic compartment during early development, including specification of the primitive endoderm from NANOG+ cells, the embryonic mesoderm lineage during gastrulation, and the primordial germ cells (5, 35). However, recent evidence demonstrated BMP activation in the TE of human, but not murine, blastocysts (5, 12), though autocrine BMP signaling has been reported in both mouse and human TSCs (36). *In vitro*, exogenous BMP4 has been shown to drive differentiation of human pluripotent stem cells (PSCs) into several lineages, namely embryonic mesoderm, primordial germ cells, or extra-embryonic trophoblast lineage progenitors, based on the cellular context and the activation of other signaling pathways (5). In particular, embryonic mesoderm differentiation requires BMP-induced WNT activation, while inhibition of β-catenin-dependent WNT signaling downstream of BMP effectively prevents mesoderm differentiation and promotes the trophoblast lineage (37). However, the mechanisms underlying WNT activation by BMP4 and the switch from mesoderm to trophoblast lineage in PSC-derived TE-like cells have not been elucidated yet.

Regulatory networks drive establishment and maintenance of cellular identity by acting on a lineage-specific epigenetic landscape downstream of signaling pathways (38). DNA methylation and histone modifications surrounding lineage-specific genes determine chromatin accessibility for specific transcription factor (TF) binding (38). Recent studies in pre-implantation embryos have described epigenome changes across early development from zygote to blastocyst, including segregation of TE and ICM (39–41). However, there is little mechanistic understanding on how epigenetic marks are laid out in a lineage-specific manner during trophoblast development. Moreover, it is largely unknown how multiple signaling pathways like HIPPO, WNT, and BMP converge towards a lineage-specific outcome and drive trophoblast development.

PSCs represent a unique resource to study species-specific genetic and epigenetic mechanisms of TE and trophoblast specification as they are tractable models for genetic and chemical manipulation. Multiple protocols have been published to derive TE from human PSCs (42–46). We have established a 2-step protocol based on BMP4 treatment of PSCs to first convert them into TE-like cells, followed by either further differentiation into mature trophoblasts or derivation of *bona fide* hTSCs (47, 48). Based on prior literature (37), we further optimized the first step by adding a WNT inhibitor, IWP2, to block concurrent activation of a mesoderm program and obtain a more homogeneous population of TE-like cells (37, 45). We previously reported that VGLL1 is one of the most highly upregulated genes in the BMP4-directed conversion of human embryonic stem cells (hESCs) into TE-like cells (9). Here, we use this model to first investigate the BMP4-dependent chromatin modifications that occur during the transition from PSC to TE-like cells in relation to gene expression changes and WNT activation status. Then, we present our findings on how VGLL1 directly and indirectly drives a trophoblast-specific gene expression and epigenomic program in this PSC-derived trophoblast model.

## Results

### BMP4 mediates chromatin accessibility to stabilize a trophoblast-specific expression program

We have previously optimized a protocol for lineage conversion of PSCs into TE-like cells by treatment with 10ng/mL BMP4 and the WNT inhibitor IWP2 over 4 days, which prevents mesoderm differentiation and promotes a uniform EGFR+ population (45). To start dissecting the transcriptional changes towards TE-like cells during this conversion protocol, we first performed bulk RNA-seq analysis of H9 hESCs at day 0, day 2, day 4 following treatment with BMP4 or BMP4/IWP2 (**Suppl. Fig. S1A**) and identified differentially expressed genes for each timepoint and treatment compared to d0 (**Suppl. Table S1**). Genes downregulated in both treatments included pluripotency and embryonic lineage markers, i.e. *NANOG*, *POU5F1*, and *SOX2* (**Suppl. Table S1**). Upregulated genes common across timepoints and treatments included the TEtra factors *GATA2*, *GATA3*, *TFAP2A*, and *TFAP2C*. By day 4, common upregulated genes included *ENPEP*, and trophoblast markers *VGLL1* and *TP63 (***Suppl. Fig. S1B and Suppl. Table S1**). Genes uniquely upregulated by BMP4 treatment on day 2 were associated with various signaling pathways, including WNT signaling (**Fig. 1A, Suppl. Table S1**) and, as expected, we detected upregulation of WNT-dependent mesoderm markers *EOMES*, *TBXT*, and *TBX6* (**Fig. 1B**). However, by day 4, the expression of these markers decreased to levels comparable to the treatment with the addition of IWP2 (**Fig. 1B**). Accordingly, WNT signaling was no longer one of the top significantly enriched pathways at day 4 of BMP4 alone treatment (**Suppl. Fig. S1C**). To confirm the activation timeframe of the WNT pathway during BMP4 treatment, we used a TCF/LEF-eGFP reporter hESC line (HUES9) where detection of GFP fluorescence is dependent on canonical WNT activation (49). We detected WNT signaling activation starting at day 2, a peak at day 3, and decreased activity at day 4 of BMP4 treatment (**Suppl. Fig. S1D**). The WNT inhibitor IWP2 completely blocked eGFP expression, while another inhibitor, IWR1, showed only partial suppression of WNT activity at the dose used (**Suppl. Fig. S1D**). Accordingly, we detected a dose-dependent inhibition of both WNT targets, *LEF1* and *WNT5B*, as well as mesoderm markers *TBXT* and *EOMES* between the two inhibitors (**Suppl. Fig. S1E**). Finally, we compared our data to the human embryo reference created by Zhao *et al.* (50) (**Suppl. Fig. S1F-G**). As expected, undifferentiated hESCs at d0 were annotated as Epiblast (**Suppl. Fig. S1F-G** and **Suppl. Table S2**). Although treatment with BMP4 and

**Figure 1.**
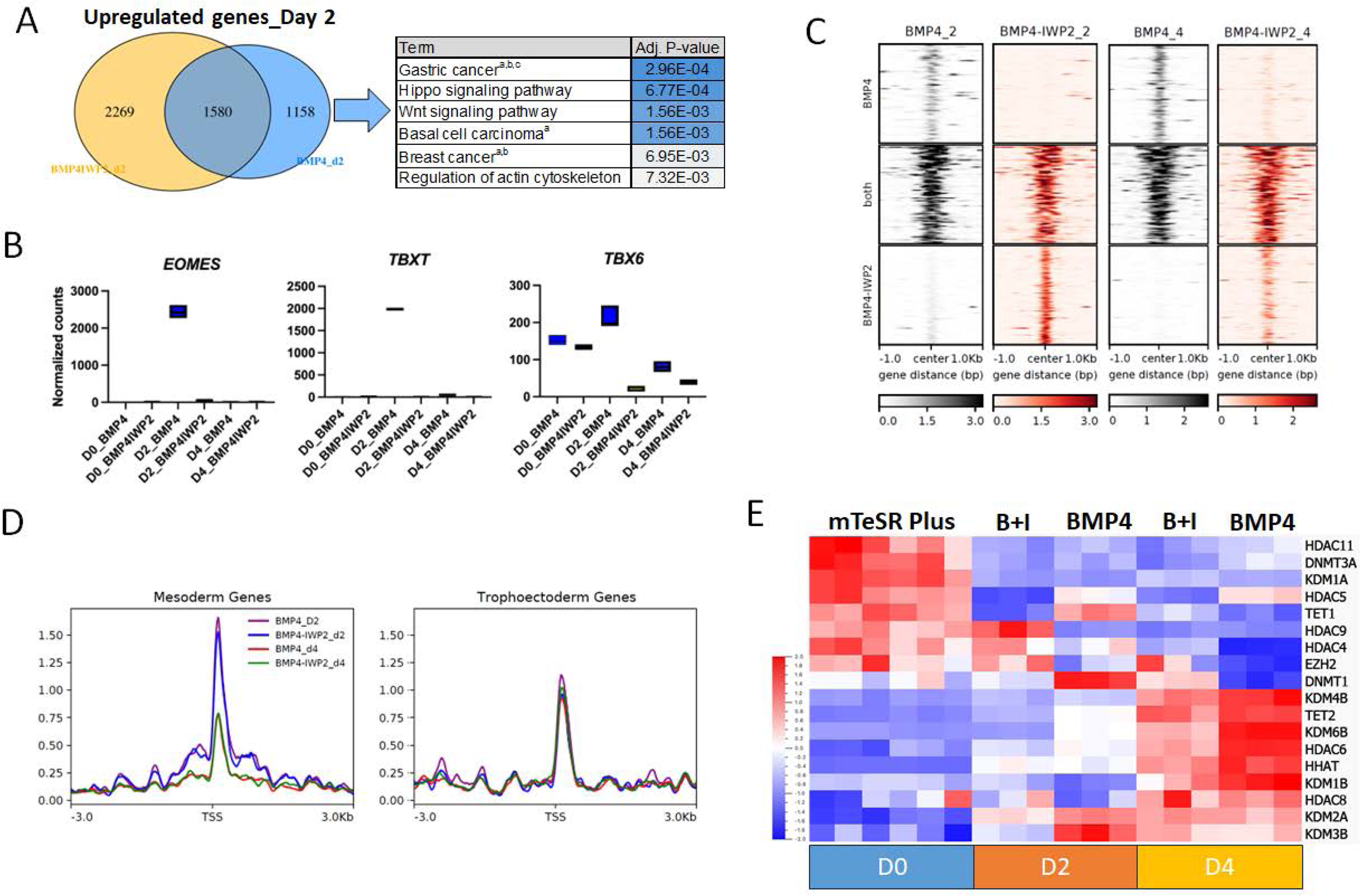
BMP4 treatment of hESCs induces chromatin remodeling. **A.** Venn diagram of up-regulated genes following treatment of hESCs with BMP4 with and without IWP2 for 2 days. Gene list enrichment analysis on the KEGG pathway database (2021) for genes uniquely up-regulated with BMP4 alone treatment is presented, a=WNT-mediated b=FGF-mediated c=TGFβ-mediated. **B.** Gene expression levels of markers for the mesoderm lineage across treatment/days by bulk RNA-seq. **C.** Heatmaps illustrating open chromatin loci by ATAC-seq. **D.** Signal intensity of open chromatin at mesoderm- and TE-specific genes. **E.** Heatmap of differentially expressed chromatin modifiers genes across treatment/days.

BMP4/IWP2 for 2 days did not sufficiently modify the transcriptome, the annotation predictor scores were lower than for hESCs at d0 (**Suppl. Fig. S1F-G** and **Suppl. Table S2**). BMP4/IWP2-treated cells at d4 were annotated as TE but only one of the BMP4 d4 replicates was also identified as TE, albeit with a lower predictor score, while the other replicates remained ambiguous and mapped close to the hypoblast (**Suppl. Fig. S1F-G** and **Suppl. Table S2**). Overall, these data suggest that low dose of BMP4 transiently activates WNT signaling during treatment of hESCs, but that both WNT activity and mesoderm gene expression decrease over time as the TE identity is established in this model.

To investigate chromatin accessibility changes associated with the differential establishment of the mesoderm and TE lineage during BMP4 treatment of PSCs, we performed Assay for Transposase-Accessible Chromatin (ATAC) followed by sequencing in hESCs treated with BMP4 or BMP4/IWP2 at days 0, 2, and 4. We found that peaks resided at promoter (31%), intergenic (24%), and intronic (23%) regions (**Suppl. Fig. S1H**). Following differential peak analysis, both at day 2 and day 4 of treatment, we found open chromatin regions specific to BMP4 or BMP4/IWP2, as well as regions common to both treatments (**Fig. 1C, Suppl. Table S3**). We performed Hypergeometric Optimization of Motif EnRichment (HOMER) analysis on common and day/treatment-specific regions to identify accessible TF binding motifs. Common regions at both days 2 and 4 contained binding motifs for well-known TE and trophoblast TFs, including TEAD and AP1-complex members (i.e. JUN and FOS) (**Suppl. Fig S1I**). Binding motifs of both TEtra factors GATA3 and TFAP2C appeared in both common and treatment-specific regions. At day 2 of BMP4-only treatment, we found open chromatin at LEF-1 and TCF3, TFs downstream of the WNT pathway. We then looked at the chromatin regions around the transcription start sites of well-known trophoblast and mesoderm genes (**Fig.1D, Suppl. Fig. S1J**). The open chromatin region across treatments/days showed a sharp peak and remained consistent around trophoblast markers (**Fig. 1D**), including GATA3 and EGFR (**Suppl. Fig. S1J**). Conversely, open chromatin around mesoderm genes showed a broader distribution that decreased between day 2 and day 4 in response to both treatments (**Fig. 1D**), including at EOMES and TBXT (**Suppl. Fig. S1J**) These chromatin accessibility data are largely consistent with the gene expression patterns observed above.

Finally, we looked at the differential expression of known epigenetic modifiers that could account for these changes in chromatin accessibility. We found multiple lineage-specific chromatin modifiers differentially expressed during TE-specification. These included down-regulation of *de novo* DNA methyltransferase *DNMT3A* at day 2, a switch from *TET1* to *TET2* (51) expression, and up-regulation of histone demethylases like *KDM6B*, *KDM4B,* and *KDM1B* (**Fig. 1E**). These data suggest that low levels of BMP4 preferentially enable a trophoblast transcriptional signature via a combination of lineage-specific gene expression activation and finely tuned regulation of the epigenetic landscape.

### VGLL1 regulates a TE-specific gene expression program

In the context of BMP4 treatment of human PSCs, the TEtra factors GATA2, GATA3, TFAP2A, and TFAP2C have been identified as initiators of the TE/trophoblast transcriptional program, upstream of more mature trophoblast markers like TP63, EGFR, ELF5, and VGLL1(9, 19, 20). To further tease apart the timeframe of activation of these TE/trophoblast genes in this BMP4-based model, we investigated the expression of TEtra factors, and mature trophoblast markers by qPCR at short intervals within the first 72 hours of BMP4/IWP2 treatment. As previously reported, TEtra factors *GATA3* and *TFAP2C* were the first to be activated, with significant upregulation as early as 6h after the start of the treatment and continuous increase over time (**Fig. 2A**). *VGLL1* expression started to significantly rise at 24 hours of BMP4/IWP2 treatment (**Fig. 2A**) while other mature TE/trophoblast markers were observed at 48 hours (*TP63*) and 72 hours (*ELF5* and *EGFR*), respectively (**Fig. 2A**). By immunofluorescence, VGLL1 protein was detected in the nucleus after 24 hours of BMP4/IWP2 treatment, where it co-localized with TEAD4 and YAP1 throughout TE induction (**Suppl. Fig. S2A**). While previous literature considered VGLL1 a mature trophoblast marker (like TP63), these temporal activation data suggest that VGLL1 might play an early role in TE induction downstream of TEtra factors.

**Figure 2.**
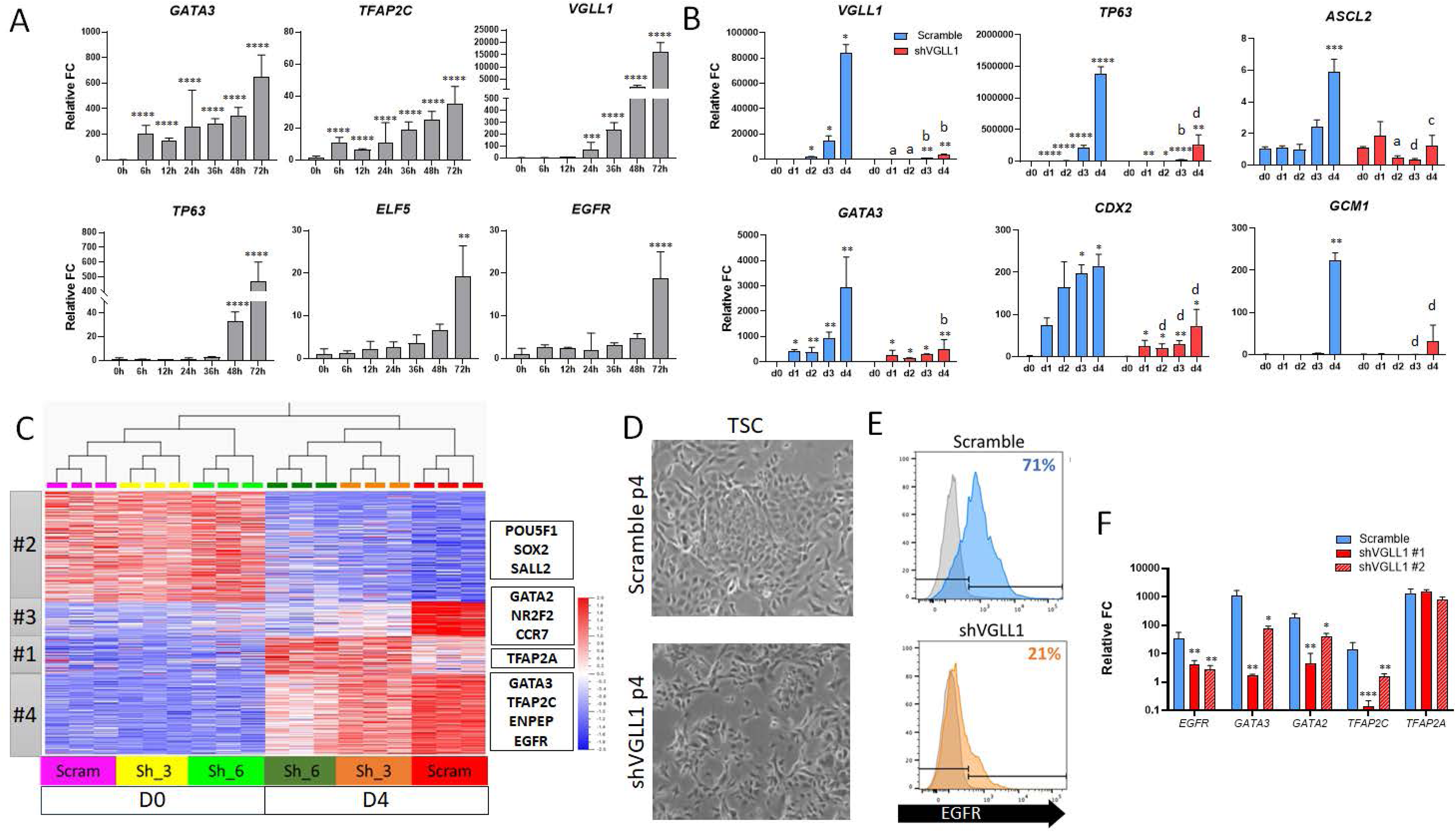
Loss of VGLL1 prevents completion of the TE/trophoblast program. **A.** Gene expression levels of early and late trophoblast markers in the first 72h of BMP4/IWP2 treatment of hESCs by qPCR. Statistical analysis relative to 0h **B.** Gene expression levels of early and late trophoblast markers in shVGLL1 compared to Scramble hESCs during BMP4/IWP2 treatment by qPCR. Asterisks represent statistical differences within cell lines compared to d0. Letters represent statistical significance of shVGLL1 compared to same timepoint on Scramble. **C.** Heatmaps of differentially expressed genes between shVGLL1 and Scramble hESCs at d0 and d4 of BMP4/IWP2 treatment by RNA-seq organized by gene cluster analysis. **D.** Bright field images of Scramble and shVGLL1 hESC-derived TSCs. **E.** Cell surface expression of EGFR in hESC-derived TSCs by flow cytometry. **F.** Gene expression levels of TEtra factors and EGFR in hESC-derived TSCs by qPCR. Statistical analysis compared to Scramble. * p≤0.05, ** p≤0.01, *** p≤0.001, **** p≤0.0001.

To understand the role of VGLL1 in this context, we established hESC lines that constitutively express VGLL1-specific shRNA for targeted gene knock-down (or shScramble as control, hereafter referred to as Scramble). Compared to Scramble hESCs, treatment of shVGLL1 hESCs with BMP4/IWP2 led to blunted up-regulation of late trophoblast markers, *TP63*, *ASCL2*, and *GCM1* (**Fig. 2B**). Interestingly, TEtra factor *GATA3* and the early TE marker *CDX2* also showed lower up-regulation. In addition, surface levels of EGFR levels were also lower in the BMP4/IWP2-treated shVGLL1-hESCs, compared to similarly-treated Scramble-hESCs (**Suppl. Fig. S2B**), suggesting an incomplete TE conversion. To further explore the effect of VGLL1 knock-down in TE induction, we performed bulk RNA-seq analysis on two independent shVGLL1 clones and one Scramble control at days 0 and 4 of BMP4/IWP2 treatment (**Suppl. Fig. S2C**). Overall, shVGLL1 hESC clones showed a marked delay in lineage conversion compared to Scramble hESCs (**Suppl. Fig. S2C**). We found 7,284 genes that were differentially dysregulated in BMP4/IWP2-treated shVGLL1 hESCs compared to Scramble hESCs (adj. p value<0.05), organized in 4 distinct clusters based on pattern of expression (**Suppl. Table S4**). Specifically, 3,056 genes showed blunted down-regulation (cluster #2) compared to Scramble and were associated with transcriptional regulation of pluripotent stem cells and embryo development (**Suppl. Fig. S2D-E**), including pluripotency markers *POU5F1*, *SOX2*, and *SALL2*. The 4,192 up-regulated genes that showed misregulation in shVGLL1 hESCs compared to Scramble hESCs (**Fig. 2C**) organized in 3 clusters of expression (**Suppl. Fig. S2D-E**). Cluster #3 contained genes that failed to be up-regulated, including TE/trophoblast markers *GATA2*, *NR2F2* (52), and *CCR7* (53), while cluster #4 included genes with blunted expression compared to Scramble, including TEtra factors *GATA3* and *TFAP2C*, TE markers *ENPEP* (54), and trophoblast markers *EGFR*, *ARID3A*, and *ARID5B* (55). Gene ontology analysis indicated that genes that failed to be fully up-regulated were associated with response to hormones and cell migration and were highly enriched for placental genes (**Suppl. Fig. S2F**). Interestingly, the remaining cluster #1 contained genes that were more highly up-regulated in shVGLL1 hESCs compared to Scramble hESCs and included TFAP2A (**Suppl. Fig. S2D-E**), a potential compensatory response to lack of VGLL1 and lower expression of other TEtra factors. Finally, we compared these cells to the human embryo reference in Zhao *et al.* (50) (**Suppl. Fig. S2G**). As expected, both Scramble and shVGLL1 undifferentiated hESCs (d0) were annotated as Epiblast (**Suppl. Fig. S2G** and **Suppl. Table S2**). BMP4/IWP2 treated cells were annotated as TE; however, Scramble and shVGLL1 cells mapped to different areas of the reference dataset, with Scramble representing more mature TE compared to shVGLL1 cells (**Suppl. Fig. S2G** and **Suppl. Table S2**). These data suggest VGLL1 acts through a positive feedback mechanism downstream of the TEtra factors to complete a TE/trophoblast-specific program.

We then investigated if VGLL1 was required for TSC derivation. Recently, we demonstrated that hESC-derived TE-like cells (hESCs following 4 days of BMP4/IWP2 treatment) can adapt to hTSC culture conditions, resulting in cells that show characteristics of *bona fide* primary hTSCs (48). hTSC culture medium includes a canonical WNT agonist and EGF signaling activation combined with TGFβ and ROCK signaling inhibition. Therefore, we attempted to derive hTSC-like cells from both Scramble and two shVGLL1 hESC clones. Similar to our initial report with wild type PSCs, both Scramble and shVGLL1 hESCs rapidly lost both epithelial morphology and cell surface expression of EGFR during the early stages of adaptation (**Suppl. Fig. S3A-B**). However, while Scramble cells acquired a hTSC-like colony morphology and regained expression of surface EGFR by passage 6, shVGLL1 clones maintained a single cell morphology and failed to regain EGFR expression (**Fig. 2D-E**). Moreover, shVGLL1 clones expressed lower levels of TEtra factors *GATA2*, *GATA3*, and *TFAP2C*, compared to Scramble hTSCs (**Fig. 2F, Suppl. Fig. S3D**). Interestingly, while the TE marker *ENPEP* also decreased in shVGLL1 cells, *CDX2* increased compared to Scramble (**Suppl. Fig. S3C-D**). We also tested the ability of TSC-adapted shVGLL1 cells to differentiate into STB and EVT following established protocols (31, 48). Unlike Scramble TSCs, TSC-adapted shVGLL1 cells failed to fuse following STB differentiation and did not secrete hCG into the media (**Suppl. Fig. S3E-F**). In EVT differentiation media, TSC-adapted shVGLL1 cells showed no morphological changes and failed to up-regulate HLA-G (**Suppl. Fig. S3G-H**). These data suggest that VGLL1 knock-down PSCs failed to form *bona fide* TSCs and that VGLL1 is required for the maintenance of a trophoblast signature, in particular related to the expression of some TEtra factors and EGFR.

We further evaluated VGLL1 activity, uncoupled from BMP4 treatment, to investigate the function of VGLL1 in promoting the TE-specific transcriptional program. We leveraged a system composed of a doxycycline (DOX)-inducible dCas9-VPR and VGLL1-specific guide RNAs for CRISPR-mediated gene activation in undifferentiated hESCs (iVGLL1a) (**Fig. 3A**). In the absence of exogenous BMP4, ectopic expression of VGLL1 in hESCs initiated changes in cell morphology towards a cobblestone-like shape (**Fig. 3B**), and increased surface expression of EGFR (**Fig. 3C**). Moreover, we observed up-regulation of the TEtra factor *GATA3* and trophoblast marker *EGFR* accompanied by down-regulation of pluripotency marker *NANOG* by qPCR (**Fig. 3D**). We then performed bulk RNA-seq analysis on three independent iVGLL1a clones treated with DOX (or DMSO as vehicle control) (**Suppl. Fig. S4A**). We found 6,530 differentially expressed genes (DEGs) in DOX-treated cells compared to DMSO (adj. p-value 0.05), of which 3,053 were up-regulated and 3,477 down-regulated (**Fig. 3E, Suppl. Table S5**). Significantly down-regulated genes were enriched in pluripotency genes, including *MYC*, *NANOG*, *POU5F1*, and *SOX2* (**Fig. 3E and Suppl. Fig. S4B**). Up-regulated DEGs included TE/trophoblast markers *ARID5B*, *EGFR*, and *GATA3* (**Fig. 3E**). Moreover, gene ontology analysis of pathways and biological processes revealed the activation of signaling pathways important for TE and trophoblast specification and function, including TGFβ, MAPK, and PI3K-Akt pathways (**Suppl. Fig. S4C**). Interestingly, DOX-treated iVGLL1a cells did not transform into TSCs following culture in TSC media: their morphology quickly reverted to compact cells and were not distinguishable from DMSO-treated cells in TSC media (**Suppl. Fig. S4D**). DOX-treated cells lost expression of VGLL1 when passaged into TSC media without DOX induction (**Suppl. Fig. S4E**). They also lost expression of the trophoblast marker TP63 as determined by qPCR (**Suppl. Fig. S4E**) and cell surface expression of EGFR as determined by flow cytometry. Interestingly, both DMSO and DOX-treated cells in TSC media showed low expression levels of the pluripotency marker *NANOG* (**Suppl. Fig. S4E**) without showing signs of differentiation into any other embryonic lineage. These data point to a specific role of VGLL1 in controlling a TE/trophoblast transcriptional program but demonstrated that ectopic expression of VGLL1 alone is insufficient for TSC establishment from hPSCs in this model.

**Figure 3.**
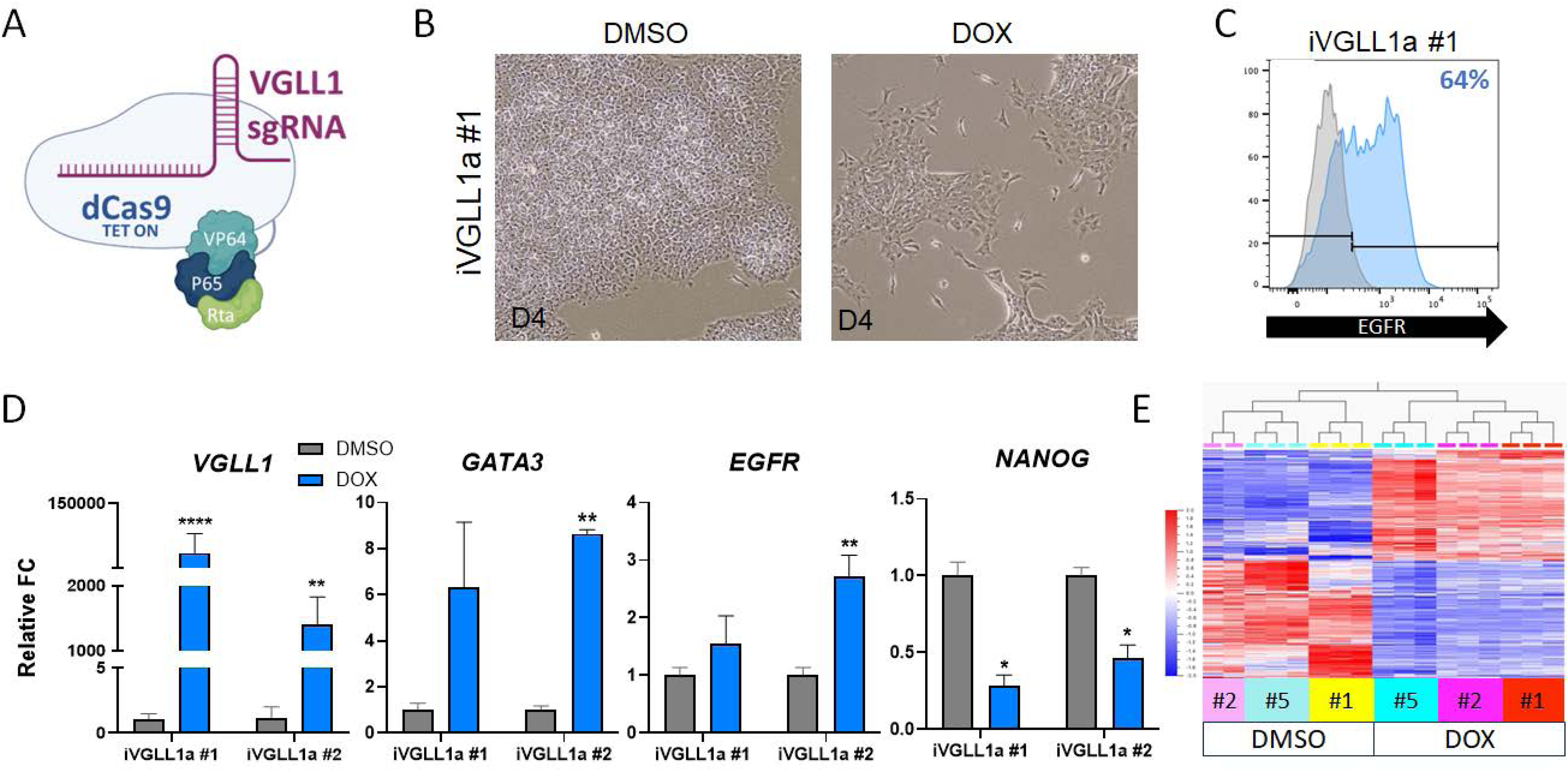
VGLL1 regulates a TE/trophoblast program independently of BMP4 treatment. **A.** Schematic representation of the inducible CRISPR-mediated gene activation system, which includes a doxycycline (DOX)-inducible dCas9-VPR and VGLL1-specific guide RNAs. **B.** Bright field images of DMSO (control) and DOX-treated hESCs at day 4. **C.** Cell surface expression of EGFR in DOX-treated iVGLL1a hESCs at day 4 by flow cytometry. **D.** Gene expression levels of lineage markers in DOX-treated iVGLL1a hESCs at day 4 by qPCR. Statistical analysis compared to DMSO. **E.** Heatmap of differentially expressed genes in iVGLL1a hESCs at day 4 of DMSO or DOX treatment. * p≤0.05, ** p≤0.01, *** p≤0.001, **** p≤0.0001

Unlike primed hPSCs, which requires BMP4, naïve hPSCs can be readily converted into TSCs following culture into TSC media (56). Therefore, we also tested the effect of ectopic VGLL1 expression in naïve hESCs on activation of TE-specific genes. We converted primed iVGLL1a hESCs to the naïve state in PXGL media (**Suppl. Fig. S4F-G**) (43). Interestingly, we observed higher basal expression of some TE/trophoblast markers in naïve hESCs compared to the primed state, including *VGLL1* and *GATA3* (**Suppl. Fig. S4G**). Similar to what we observed in primed hESCs, treatment with DOX and consequent VGLL1 up-regulation caused up-regulation of TE markers *GATA3* and *EGFR* as well cell surface expression of EGFR (**Suppl. Fig. S4H-I**). These data demonstrate that the program initiated by VGLL1 is conserved independently of pluripotency state.

### VGLL1 regulates canonical WNT pathway effectors

Gene ontology analysis of up-regulated genes in response to DOX treatment of iVGLL1a hESCs suggested activation of WNT signaling pathway (**Suppl. Fig. S4C**). Indeed, VGLL1 expression induced up-regulation of WNT pathway components and mediators, including *EPHA2*, *FZD2*, *LEF1*, *LRP4*, *WLS*, *WNT4/5A/5B/6/10A/11*, as well as WNT signaling targets like *FST* (**Fig. 4A and Suppl. Fig. S4J**). We also confirmed b-catenin dependent WNT activation using a reporter expressing mCherry under constitutive promoter and GFP under TCF/LEF promoter (57). We observed GFP signal in a subset of mCherry expressing cells following DOX treatment compared to DMSO control (**Fig. 4B**). Interestingly, we observed up-regulation of mesoderm markers on DOX-treated iVGLL1a hESCs by qPCR, including *EOMES* and *TBXT* (**Fig. 4C**). During BMP4-induced hESC differentiation, co-treatment with WNT inhibitor IWP2 effectively suppressed mesoderm marker expression. Therefore, we tested the dependence of mesoderm gene expression down-stream of VGLL1 activation in iVGLL1a hESCs on activation of the WNT signaling pathway. Addition of IWP2 during doxycycline induction of VGLL1 effectively reduced the expression of WNT genes *FST*, *LEF1*, and *LGR5* (**Suppl. Fig. S4JH**). Moreover, this resulted in reduced expression of mesoderm markers *EOMES* and *TBXT* (**Fig. 4C**) and improved expression levels of *GATA3* and *EGFR* by qPCR and flow cytometry (**Fig. 4D-E**). We compared gene expression profiles of DOX and DOX/IWP2 treated cells (**Fig. 4F, Suppl. Table S6**) and found that DOX/IWP2 treated cells showed significant loss of mesoderm-associated genes, including *EOMES* and *TBXT* (**Suppl. Table S6**). Interestingly, ectopic expression of VGLL1 in naïve hESCs showed similar up-regulation of WNT-related genes but not mesoderm markers, *EOMES* and *TBXT* (**Suppl. Fig. S4L**). These data support a role for VGLL1 in regulating the expression of β-catenin-dependent WNT signaling genes, an important pathway in trophoblast development and placental function.

**Figure 4.**
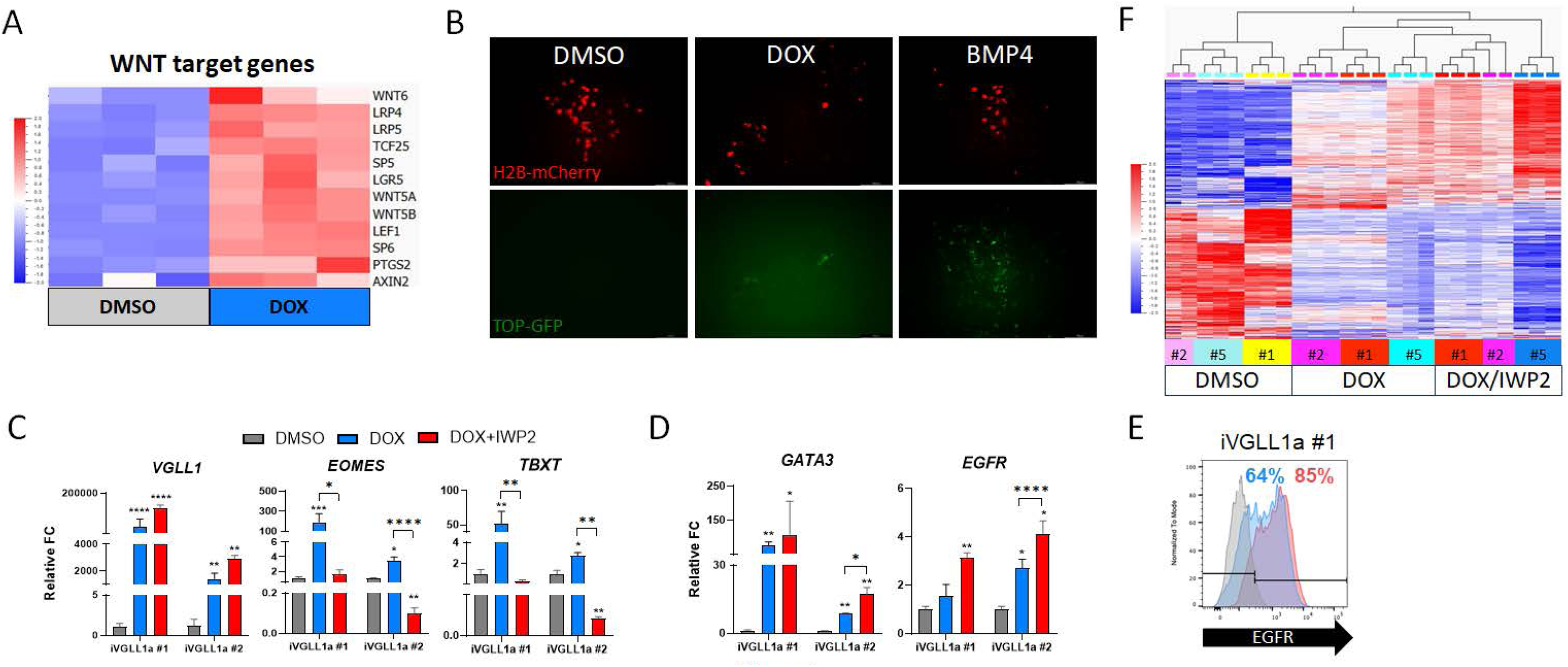
VGLL1 regulates expression of canonical WNT effectors. **A.** Heatmap of WNT target genes differentially expressed in iVGLL1a hESCs at day 4 of DMSO or DOX treatment. **B.** β-catenin-dependent WNT activation in DOX-treated hESC using a TCF/LEF-GFP reporter plasmid. **C.** Gene expression levels of mesoderm markers in DOX-treated iVGLL1a hESCs with and without IWP2 by qPCR. **D.** Gene expression levels of early and late trophoblast markers in DOX-treated iVGLL1a hESCs with and without IWP2 by qPCR. **E.** Cell surface expression of EGFR in DOX-treated iVGLL1a hESCs with and without IWP2 at day 4 by flow cytometry. **F.** Heatmap of differentially expressed genes in iVGLL1a hESCs at day 4 of DMSO or DOX treatment, with and without IWP2. * p≤0.05, ** p≤0.01, *** p≤0.001, **** p≤0.0001.

### VGLL1 directly regulates trophoblast-associated genes at placenta-specific distal regulatory regions

We compared the differentially expressed genes in both the constitutive knock-down and gene activation systems to further narrow down VGLL1 contributions to the TE transcriptional program (**Fig. 5A**). We found 528 genes that were concordantly regulated by VGLL1 expression, specifically down-regulated in VGLL1sh hESCs (relative to Scramble) after TE induction, and up-regulated in the iVGLL1a hESCs treated with DOX/IWP2 (**Suppl. Table S5**). These genes were involved in cancer pathways, focal adhesion, TGFβ and HIPPO signaling pathways (**Suppl. Fig. S4E**) and were highly associated with placental gene expression (**Suppl. Fig. S5A**). We then set out to investigate whether VGLL1 directly regulates the transcription of a subset of these genes. VGLL1 lacks a DNA binding domain but is a transcriptional co-factor for TEAD family members. TEAD4 is expressed in both hESCs and hESC-derived TE-like cells and plays a role in trophoblast differentiation of hESCs as well as in maintenance of stemness in primary hTSCs (58, 59). Therefore, we predicted that VGLL1 binding to regulatory regions might be mediated through its association with TEAD4. To test this, we performed immunoprecipitation with either an anti-TEAD4 or anti-VGLL1 antibody on the choriocarcinoma cell line JEG3 and confirmed co-precipitation of VGLL1 and TEAD4, respectively (**Suppl. Fig. S5B**). We further confirmed VGLL1 binding to TEAD4 in BMP4-IWP2-treated TE-like cells by co-immunoprecipitation (**Suppl. Fig. S5B**). We also confirmed binding of VGLL1 and TEAD4 to a validated 5’UTR region of *GATA3* by ChIP-qPCR (**Suppl. Fig. S5C**) (60).

**Figure 5.**
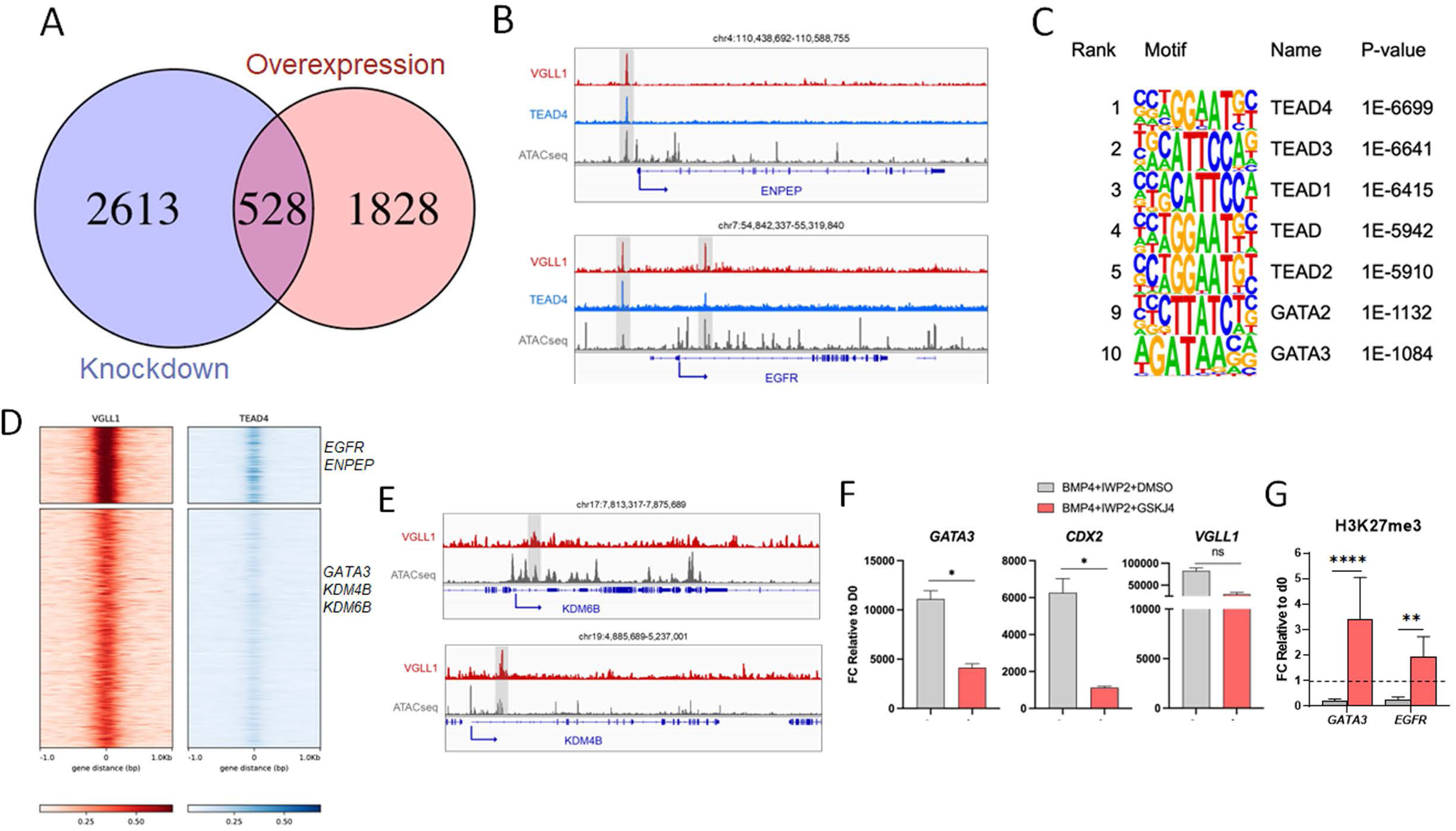
VGLL1 directly regulates chromatin modifier gene KDM6B. **A.** Venn diagram of concordantly regulated genes downregulated in shVGLL1 clones and up-regulated in DOX-treated iVGLL1a hESCs. **B.** Genome browser tracks showing VGLL1 and TEAD4 genomic enrichment peaks (ChIP-seq) and chromatin accessibility peaks (ATAC-seq) for representative TE gene loci of BMP4/IWP2-treated hESCs at day 4. **C.** Motif analysis using HOMER showing the significantly enriched DNA-binding motifs at VGLL1-binding sites in BMP4/IWP2-treated hESCs at day 4. **D.** Co-occupancy analysis by signal density pileups of VGLL1 and TEAD4 peaks in ChIP-seq of BMP4/IWP2-treated hESCs at day 4. **E.** Genome browser tracks showing VGLL1 genomic enrichment peaks (ChIP-seq) and chromatin accessibility peaks (ATAC-seq) for chromatin modifiers, KDM6B and KDM4B loci of BMP4/IWP2-treated hESCs at day 4. **F.** Gene expression levels of early trophoblast markers after 4 days of treatment of hESCs with BMP4/IWP2 in the presence of KDM6B inhibitor GSKJ4 (or DMSO as control). **G.** CUT&RUN for repressive histone mark H3K27me3 followed by qPCR for regulatory regions occupied by VGLL1 of TE markers GATA3 and EGFR. * p≤0.05, ** p≤0.01, **** p≤0.0001, ns = non-significant

Having validated the interaction between VGLL1 and TEAD4, we next set out to identify binding regions of VGLL1 in complex with TEAD4 in hESC-derived TE-like cells. Therefore, we performed ChIP-seq using both anti-VGLL1 and anti-TEAD4 antibodies on BMP4/IWP2-treated hESCs at day 4. VGLL1 ChIP peaks were distributed mainly between intronic (44.4%), intergenic (30.6%), and promoter (20%) regions (**Suppl. Fig S5D**), suggesting that VGLL1 might regulate target genes through transcription activation at promoters as well as through interactions at distal enhancer regions. In line with this, we found evidence of VGLL1 occupancy at *EGFR* and *ENPEP* promoter regions (**Fig. 5B)**, as well as in distal regulatory regions of *GATA3* and *EGFR* identified as putative active enhancers in the human placenta (**Suppl. Fig. 5E, Suppl. Table S6**) (61). VGLL1 peaks were enriched in TEAD family member motifs, including TEAD4 and TEAD1 (**Fig. 5C**). In our ChIP-seq data, VGLL1 and TEAD4 co-occupied regions at trophoblast-associated genes *EGFR*, *ARID3A*, *CDH1*, *ENPEP*, *KRT7*, *KRT8*, *PTPN14*, and *TEAD1* (**Fig. 5D**, **Suppl. Fig. S5F, Suppl. Table S6**).

In addition, we identified VGLL1-specific binding at TE/trophoblast regulators *ASCL2*, *ARID5B*, *CCR7*, CITED2, *NODAL*, *TP63,* and WNT signaling effectors, such as the long non-coding *LEF-AS1*, and components *EPHA2*, *FZD2*, *ITGB1, LGR4*, *LRP5/6*, *WLS*, *WNT5A*, and *WNT5B* (**Fig. 5D and Suppl. Table S6**). The genes bound by VGLL1 were highly associated with the human TE transcriptional signature and not with mesoderm-derived cell types (**Suppl. Fig. S5G**). To confirm that these regulatory regions are specific for VGLL1 in the TE context independently of the model used, we mined published VGLL1 CUT&TAG data from Yang *et al.* in naïve TE cells (62) and found conserved binding regions in TE-specific genes, including at *GATA3*, *EGFR*, and *ENPEP loci* (**Suppl. Fig. S5H**). Together, these data demonstrate that VGLL1 directly regulates a TE/trophoblast-specific program at promoter regions in complex with TEAD4 and at distal regulatory regions potentially with a novel partner.

### VGLL1 contributes to BMP4-dependent epigenetic changes by regulating the histone demethylation enzyme KDM6B

To investigate if VGLL1 might contribute to the epigenetic changes for TE induction of hESCs downstream of BMP4-treatment, we first looked at chromatin modifiers whose expression patterns followed VGLL1 manipulation (**Fig. 5A, Suppl. Table S5**). Among the 528 genes concordantly regulated, we identified *TET2*, involved in DNA demethylation, and *KDM6B*, an enzyme responsible for demethylation of histone 3 lysine residue 27 (H3K27me3/2/1). Furthermore, we detected VGLL1 occupancy at *KDM4B* and *KDM6B* loci in BMP4/IWP2-treated hESCs (**Fig. 5D**), indicating these lysine demethylases are direct targets of VGLL1 in our model. Interestingly, we found overlapping binding regions at *KDM6B* loci in the VGLL1 CUT&TAG dataset in naïve TE cells from Yang *et al.* (62), suggesting a conserved mechanism (**Suppl. Fig. S5H**). KDM6B has been reported to regulate the demethylation of H3K27me3 at promoter regions of *Gata3* and *Cdx2*, thus promoting their expression in the TE of pre-implantation murine embryos (63). Therefore, we treated hESCs with a KDM6B demethylase activity inhibitor, GSKJ4, during BMP4/IWP2-mediated differentiation, and, similar to VGLL1 knock-down, we found decreased levels of TE markers *GATA3* and *CDX2*, compared to vehicle control (**Fig. 5F**). To confirm maintenance of the H3K27me3 repressive mark at VGLL1 binding peaks of TE-specific genes, we performed CUT&RUN with an anti-H3K27me3 antibody followed by qPCR (64). We found decreased H3K27me3 binding at both *GATA3* and *EGFR* regions occupied by VGLL1 following 4 days of BMP4/IWP2 treatment compared to d0 in control samples (**Fig. 5G**). In contrast, H3K27me3 levels at these *loci* were maintained or even increased at d4 compared to d0 upon addition of GSKJ4 (**Fig. 5G**). Collectively, these data suggest that VGLL1 further contributes to the establishment of a TE/trophoblast transcriptional signature by regulating the expression of chromatin remodeler enzymes.

### VGLL1 and KDM6B are co-expressed in the TE compartment of pre-implantation embryos

To extend our findings beyond PSC-based models, we examined VGLL1 expression in human embryos. Using published datasets, we investigated its temporal and spatial expression during early embryonic development. Yan *et al*. previously performed RNA sequencing (RNA-seq) of individual early human embryos from the oocyte to pre-implantation blastocyst (27). Analysis of this data revealed that *VGLL1* was expressed at very low levels until the blastocyst stage, at which point its expression began to increase (**Suppl. Fig. S6A**). Blakeley *et al.* investigated the transcriptional signature of the three main cell types found in the pre-implantation blastocyst: TE, epiblast (EPI), and primitive endoderm (PE) with single-cell RNA-seq (3). Examination of datasets from these cell types revealed that *VGLL1* was highly expressed in TE, with some expression in the PE but not in EPI (**Suppl. Fig. S6B**). To further distinguish between polar TE (adjacent to the inner cell mass) and mural TE (adjacent to the blastocoele) clusters, we re-analyzed single-cell RNA-seq data of dissociated human embryo cells from another study that performed extended embryo culture (65). We found that *VGLL1* expression was increased in TE relative to the EPI and PE, with slightly higher expression in the polar TE compared to the mural TE (**Fig 6A**). By contrast, *CDX2*, another TE marker central to the TE program in humans and mice, was similarly increased in TE compared to EPI and PE cells but was higher in the mural compared to the polar compartment (**Fig 6A**). To assess if protein expression correlated with our transcriptomic results, we co-stained pre-implantation embryos with VGLL1 and both extra-embryonic (GATA3, TP63) and embryonic (NANOG) markers. We observed VGLL1 positive cells at the morula stage in both inside and outside cells and co-staining with GATA3 in a subset of outside cells (**Suppl. Fig. S6C**). At the blastocyst stage (day 5), VGLL1 expression was higher in the nuclei of TE cells compared to ICM (**Fig 6B-C, Suppl. Fig. S6D**). These data confirm VGLL1 activity in the TE compartment of human embryos, further suggesting a role for this transcriptional co-factor in establishment/maintenance of this lineage.

**Figure 6.**
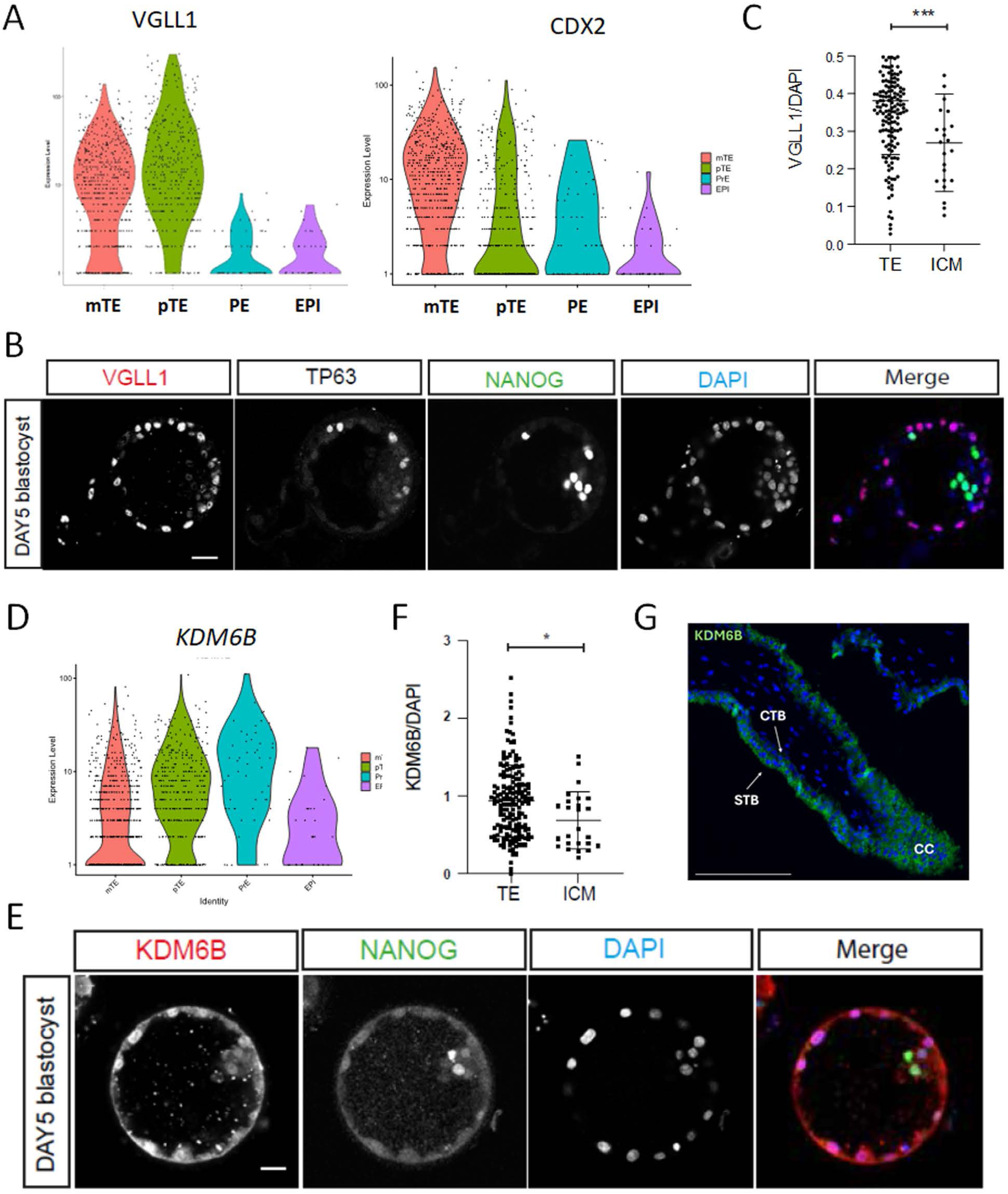
VGLL1 and KDM6B are highly expressed in the polar TE of pre-implantation blastocysts. **A.** Violin plots showing expression levels of VGLL1 and CDX2 in the mural TE, polar TE, epiblast, and primitive endoderm (PE) cells of d6 blastocysts (data from Zhou *et al.* 2019). **B.** Immunofluorescence staining of a day 5 human blastocyst with VGLL1, TP63, and NANOG. **C.** Quantification of VGLL1 staining in the TE and ICM compartments of day 5 human blastocyst (n=3) ***p=0.0005. **D.** Violin plots showing expression levels of KDM6B in the mural TE, polar TE, epiblast, and primitive endoderm (PE) cells of d6 blastocysts (data from Zhou *et al.* 2019). **E.** Immunofluorescence staining of a day 5 human blastocyst with KDM6B and TP63. **F.** Quantification of KDM6B staining in the TE and ICM compartments of day 5 human blastocyst (n=3) *p=0.0117. **G.** Immunofluorescence staining of KDM6B in a week 6 placenta. CTB= cytotrophoblast, STB=syncytiotrophoblast, CC=cell column

Finally, to determine if VGLL1 and KDM6B were found in similar compartment *in vivo*, we examined KDM6B expression in both human embryos and first trimester placentas. In RNA-seq datasets from human pre-implantation embryos, *KDM6B* was highly expressed in the TE, with increased expression in the polar compared to mural TE (**Fig. 6D**). However, it was also expressed at high levels in PE and at lower level in the ICM. In day 5 blastocysts, protein levels recapitulated observations in single cell RNA-seq datasets, with significantly higher expression of KDM6B in cells of the TE compared to those of the ICM (**Fig. 6E-F**). Finally, in the first trimester placenta, immunostaining showed ubiquitous expression of KDM6B in the trophoblast compartment with lower expression in cytotrophoblasts (CTB) compared to expression in both the proximal cell column (CC) and the syncytiotrophoblast compartment (STB) (**Fig. 6G**). Taken together with our ChIPseq, these data point to a direct role for VGLL1 in regulating KDM6B expression in the TE and the trophoblast compartment of first trimester placenta. Overall, this study demonstrates that VGLL1 regulates both genetic and epigenetic features to support a trophoblast lineage-specific transcriptional program.

## Discussion

Correct establishment of the TE lineage is fundamental for placental development and a successful pregnancy outcome. Because of the ethical and technical limitations of working with human embryos, various protocols have been reported to differentiate human pluripotent stem cells into trophoblast in order to study species-specific mechanisms of development. In this work, we used a BMP4-model for TE induction from primed PSCs previously established in our lab, which includes addition of the WNT inhibitor IWP2 to suppress mesoderm differentiation, followed by TSC culture adaptation (48). First, we further explore the transcriptional programs and lineages elicited by BMP4 in the presence and absence of IWP2. We demonstrated that activation of the canonical WNT pathway and consequent up-regulation of mesoderm markers was limited to the first 2 days of treatment, with transcription directed towards a TE-like program by day 4. Universal TE markers including *GATA3*, *ENPEP*, and *TACSTD2* were all induced by BMP4 treatment. Our ATAC-seq data comparing BMP4-driven differentiation with and without WNT signaling inhibition indicates that, in primed hPSCs, the chromatin surrounding trophoblast lineage genes was accessible for transcription throughout differentiation. The changes in chromatin reported here accompany the temporal increase of TE/trophoblast markers that we observed in our RNA-seq data. In contrast, we showed that, at these concentrations of BMP4, chromatin near mesoderm lineage genes closed over time even in the absence of WNT inhibition.

Temporal activation of TE/trophoblast lineage in our model was similar to previously published studies of BMP4-dependent induction of the trophoblast lineage, with TEtra factors GATA2/3 and TFAP2A/C expressed early during differentiation followed by more mature markers (19). However, in their study Krendl and colleagues defined VGLL1 as a late marker. In this study, we determined that *VGLL1* induction occurs after the TEtra factors but before other mature markers like *EGFR* and *TP63*. Knock-down of VGLL1 prevented a complete exit from pluripotency with maintenance of factors like *POU5F1* and *SOX2*, and blunted expression TE markers including *EGFR*, *ENPEP*, and *NR2F2*. Interestingly, we also observed reduced expression of TEtra factors *GATA3*, *GATA2*, and *TFAP2C*. These data suggest that while VGLL1 is not required for the initiation of the TE/trophoblast program, it belongs to a second wave of factors necessary for the completion of this program in part via a positive feedback loop onto the TEtra factors.

Such a role is not unique to VGLL1. In the mouse, Eomesodermin (Eomes), downstream of Cdx2, is not required to initiate the TE program (18, 66) but instead plays a key role as regulator of TE differentiation and maintenance of murine TSCs in the extra-embryonic compartment (17). Mouse embryos lacking Eomes are arrested in the early post-implantation period, initiating formation of TE (e.g. by cavitation), but failing to form a proper chorion (17). While EOMES is not involved in human TE/trophoblast development (3, 9), our data suggest that VGLL1 function similarly to reinforce the TE/TSC identity downstream of early key TE lineage regulators and allow hTSC expansion. In fact, VGLL1 knock-down TE-like cells adapted to TSC culture conditions showed decreased expression of *GATA2* and *GATA3*, lack of cell surface expression of EGFR, and a failure to differentiate into either STB or EVT cells, demonstrating lack of *bona fide* hTSC derivation.

Canonical WNT signaling is a crucial pathway to establish and maintain various epithelial stem cells (23). As mentioned, canonical WNT signaling supports mesoderm differentiation of BMP4-treated hPSCs (37). VGLL1 is uniquely expressed in the TE/trophoblast lineage during early human development, thus the up-regulation of mesoderm genes *EOMES*, *MIXL1*, and *TBXT* in hPSCs as a consequence of ectopic expression of VGLL1 was unexpected. As in our BMP4-based differentiation protocol, addition of IWP2 significantly decreased the expression of mesoderm genes in iVGLL1a hESCs, suggesting that canonical WNT activation, but not mesoderm genes, may be directly regulated by VGLL1. In fact, temporal activation of the canonical WNT reporter at 48-72h of BMP4-treatment correlated with the timeline of VGLL1 induction. We also detected active WNT signaling in DOX-treated iVGLL1a hESCs. However, our data also suggest that VGLL1 alone cannot sustain long-term autocrine activation of the canonical WNT pathway. We detected VGLL1 occupancy at WNT coreceptors *LRP5/6* as well as the WNT agonist R-Spondin receptor *LGR4*. Canonical WNT pathway activation is necessary for hTSC and trophoblast organoid establishment and maintenance from placental tissue. Our data therefore suggest that VGLL1 is key in supporting and maintaining the trophoblast stem cell state, potentially via up-regulation of canonical WNT signaling pathway receptors and coreceptors.

Recently, Yang and colleagues explored the role of VGLL1 and TEAD4 in TE specification, starting from naïve PSCs, which represent cells of the pre-implantation embryonic compartment (62). Similar to our results, they demonstrated that loss of VGLL1 led to maintenance of pluripotency genes, blunted expression of TE genes, including *GATA3*, *ENPEP* and *NR2F2*, and inability to derive hTSCs. They demonstrated that the VGLL1/TEAD4 complex directly regulates expression of markers such as *GATA3* and *ENPEP* at promoter and intergenic regions. In contrast, our model utilizes primed PSCs, which represent a post-implantation epiblast state (50). Despite the different initial states, our data largely corroborate their findings with VGLL1 directly regulating markers such as *GATA3* and *ENPEP*. Interestingly, we showed VGLL1 occupancy at putative distal regulatory regions of *GATA3* and *EGFR*. These regions were recently mapped to predicted enhancers active in the human placenta across gestation (61), suggesting a conserved role for VGLL1 in trophoblast-specific distal regulatory regions. In fact, when we looked at the presence of these distal enhancer peaks in the Yang *et al* data, we found shared peaks, corroborating that they might represent functionally relevant regulatory regions for TE and trophoblast identity.

The TE and ICM lineages segregate early during mammalian development and are distinguished by unique transcriptomes and epigenomes. Both naïve and primed hPSCs contain bivalent promoters at developmental genes, which maintain them in a transcriptionally poised state (19, 67, 68). Such bivalent regions are characterized by H3K4me3, a post-translational histone modification mark associated with active promoters, and H3K27me3, a marker correlated with transcriptional repression. Global distribution of H3K27me3 differs between primed and naive hPSCs. However, many of the genes marked by H3K27me3 that are shared between these two pluripotency states are trophoblast regulators, including GATA3 (67, 68). In their study, Yang and colleagues found a correlation between loss of VGLL1 and lower levels of H3K27ac, a histone mark associated with active promoters and enhancers, at TE-specific genes (62). However, the authors did not propose a direct link between VGLL1 expression and H3K27ac levels at TE genes. Moreover, before H3K27 can be modified by acetylation, the methyl groups need to be removed from the bivalent promoter occupied by H3K27me3. We have also observed decreased levels of global H3K27ac in shVGLL1 hESCs in our primed model of TE conversion and identified KDM6B, one of two H3K27me3-specific demethylases in mammals, as a potential mediator of VGLL1-dependent chromatin remodeling. The transcript levels of *KDM6B* were largely dependent on VGLL1 expression and we observed direct VGLL1 binding on *KDM6B* gene in our ChIP-seq data. In the murine blastocyst, KDM6B removes the repressive H3K27me3 at the promoter regions of *Gata3* and *Cdx2* specifically in the TE (63). Loss of *Kdm6b* results in maintenance of H3K27me3 at TE-specific gene regulatory regions, lower expression of Gata3 and Cdx2 in the TE, and an impaired ability for the embryos to implant into the uterus. In humans, *VGLL1* and *KDM6B* are highly expressed in the polar TE of the human blastocyst, which attaches to the uterine wall during implantation. It remains unclear whether KDM6B is involved in human implantation, but our results show that chemical inhibition of KDM6B demethylase activity phenocopies loss of VGLL1 and results in the blunted expression of *GATA3* and *CDX2*, *in vitro*. Thus, we posit a model whereby VGLL1 directly up-regulates the expression of KDM6B, which contributes to demethylation of H3K27me3 at bivalent/poised promoters prior to deposition of the acetyl group by acetyltransferases at trophoblast-specific regulatory regions to mark a transcriptionally permissive chromatin state (**Fig.7**). Evidence suggests that, while mouse TE is fully specified in the pre-implantation blastocyst, human TE lineage commitment might not be completed until post-implantation (52). VGLL1-dependent regulation of chromatin remodelers like KDM6B might provide a mechanism explaining the higher plasticity of human post-implantation pluripotent cells. In the human placenta, KDM6B was expressed across the trophoblast compartments of first trimester tissues, including villous CTB, cell column trophoblasts, and STB. Therefore, our data suggests that KDM6B may play a role not just in TE specification but also in functional mature trophoblasts.

**Figure 7.**
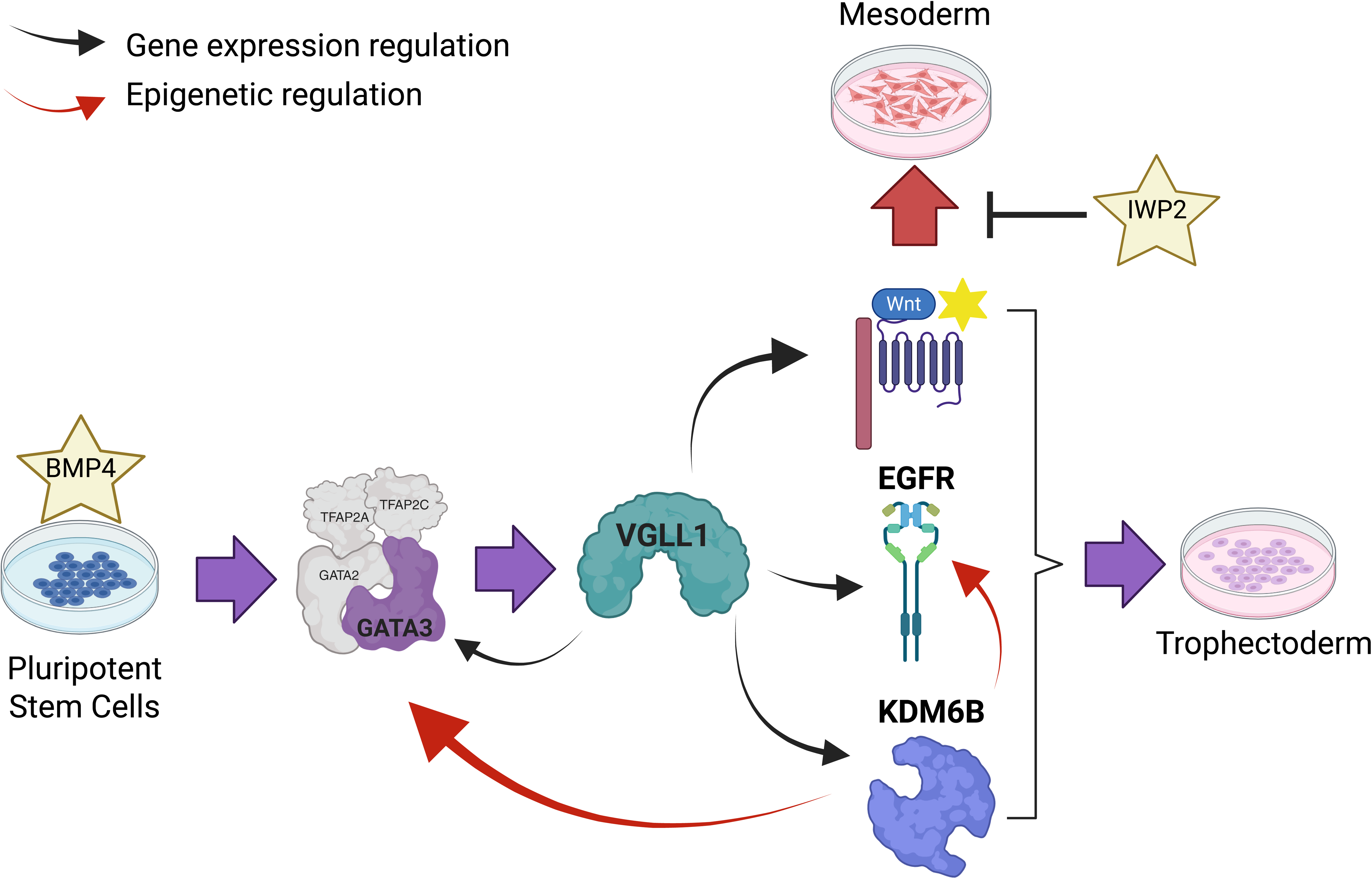
Graphic summary of the results. Created in BioRender. Soncin, F. (2025).

Overall, our data extend previous findings and add a substantial mechanistic hypothesis on the role of VGLL1 in TE and trophoblast lineage specification. We also demonstrated that primed PSCs and the BMP4-dependent conversion protocol remain a functional and practical *in vitro* model to study fundamental aspects of genetic and epigenetic regulation of TE and trophoblast cell identity in humans (69).

## Material and Methods

### Human embryo and Placental Tissue Collection

Human embryos were donated to the research project under the UK Human Fertilisation and Authority Licence number R0162. The full study protocol was approved by the Health Research Authority’s Research Ethics Committee IRAS projects 308099. Human placental tissues were collected under a protocol approved by the Human Research Protections Program Committee of the University of California, San Diego Institutional Review Board. All patients gave informed consent for the collection and use of embryos/placental tissues and samples were de-identified prior to use.

### Trophoblast differentiation of hESC

hESCs were first converted into TE-like cells followed by adaptation to hTSC culture conditions according to a previously published protocol (70). Briefly, hESCs were plated at a density of 10,000 cells per cm^2^ on Geltrex-coated plates in mTeSR Plus with 5μM Y27632 (Selleck Chemicals, cat. no. S1049) and differentiated in basal media with 10 ng/mL recombinant human BMP4 (R&D Systems, cat. no. 314-BP-010), and 2 μM IWP2 (Selleck Chemicals, cat. no. S7085). At day 4, cells were either collected for analysis or re-plated on collagen IV (5μg/mL)-coated plates (MilliporeSigma, cat. no. C0543-1VL) in iCTB media. Other reagents include 2 μM IWR1 (MilliporeSigma, cat. no. I0161) and 1 μM GSK-J4 (MedChem Express, cat. no. HY-15648B).

## Data availability

All datasets are available in Geo: GSE295121, GSE295122, GSE295124, GSE295126.

## Acknowledgments

All sequencing was conducted at the IGM Genomics Center, University of California, San Diego, La Jolla, CA. We thank Dr. Karl Willert for the generous donation of the TOPGFP HUES9 cell line, and Dr. Marisa Korody for the IWR1. We are grateful to Dr. Mana Parast for the continuous encouragement and support. The contents of this publication are solely the responsibility of the authors and do not necessarily represent the official views of CIRM or any other agency of the State of California.

