## Supplemental files for "VGLL1 contributes to both the transcriptome and epigenome of the developing trophoblast compartment"

PNAS

Francesca Soncin

**This PDF file includes:**

Supplementary Figures with legend S1 to S6
Supplementary Methods
Supplementary Tables S1-S6 titles and legends
Supplementary Table S7. Primer list
Supplementary Table S8. TE and Mesoderm genes
Supplementary References

**Other supporting materials for this manuscript include the following:**

Supplementary Tables S1-S6 (large datasets uploaded individually as excel files)

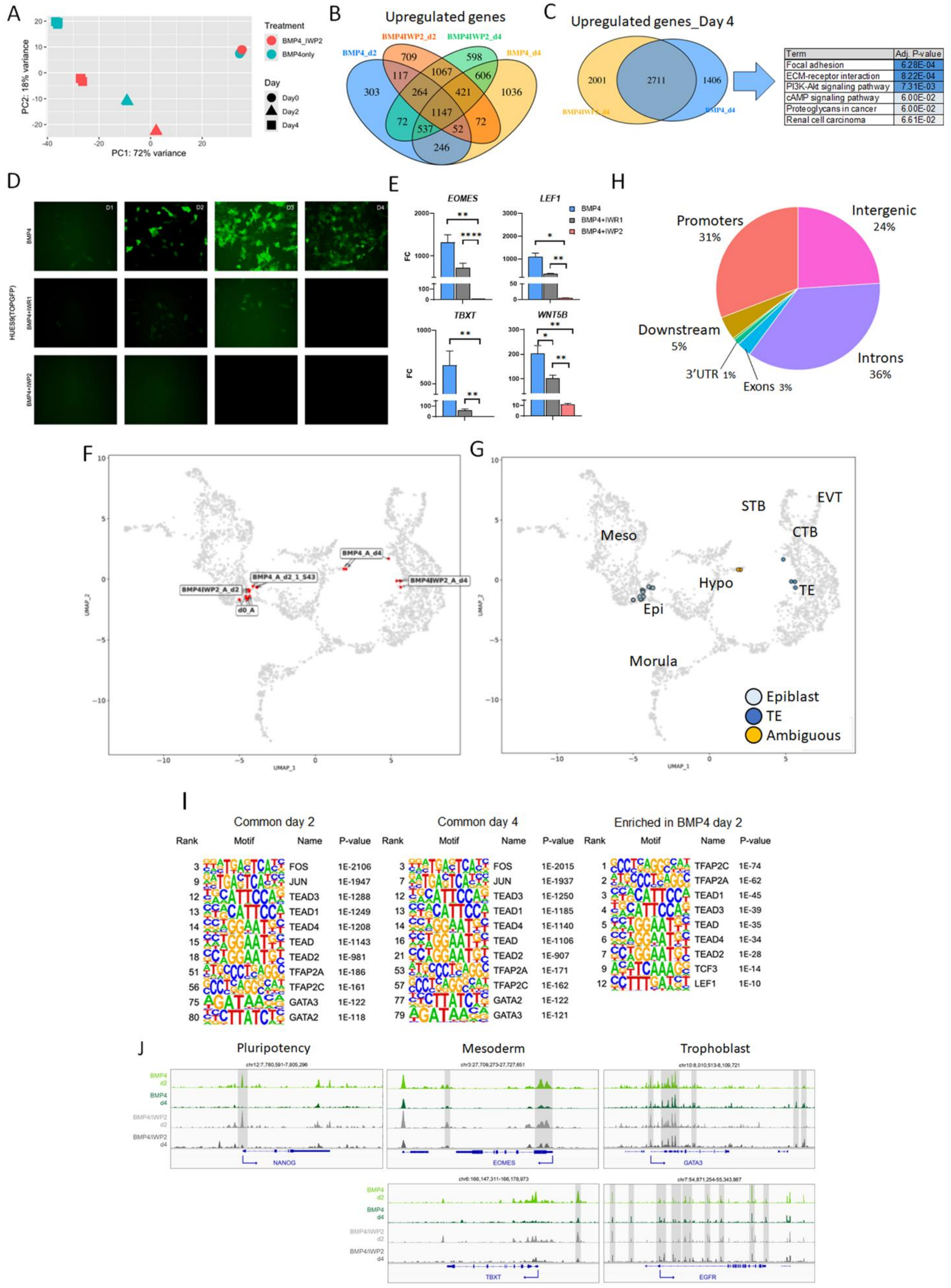
A
Treatment
BMP4_IWP2
BMP4only
Day
Day0
Day2
Day4
PC1: 72% variance
PC2: 18% variance
B
Upregulated genes
BMP4_d2
BMP4IWP2_d2
BMP4IWP2_d4
BMP4_d4
C
Upregulated genes_Day 4
2001
2711
1406
Term
Adj. P-value
Focal adhesion
6.28E-04
ECM-receptor interaction
8.22E-04
PI3K-Akt signaling pathway
7.31E-03
cAMP signaling pathway
6.00E-02
Proteoglycans in cancer
6.00E-02
Renal cell carcinoma
6.61E-02
D
BMP4
BMP4+IWR1
BMP4+IWP2
HUES9(TOPGFP)
D1
D2
D3
D4
E
EOMES
LEF1
TBXT
WNT5B
FC
BMP4
BMP4+IWR1
BMP4+IWP2
H
Promoters 31%
Intergenic 24%
Downstream 5%
3'UTR 1%
Exons 3%
Introns 36%
F
BMP4_A_d4
BMP4_A_d2_1_543
BMP4IWP2_A_d2
BMP4IWP2_A_d4
d0_A
UMAP_1
UMAP_2
G
STB
EVT
CTB
Meso
Hypo
TE
Epi
Morula
Epiblast
TE
Ambiguous
I
Common day 2
Common day 4
Enriched in BMP4 day 2
Rank
Motif
Name
P-value
3 FOS 1E-2106
9 JUN 1E-1947
12 TEAD3 1E-1288
13 TEAD1 1E-1249
14 TEAD4 1E-1208
15 TEAD 1E-1143
18 TEAD2 1E-981
51 TFAP2A 1E-186
56 TFAP2C 1E-161
75 GATA3 1E-122
80 GATA2 1E-118
3 FOS 1E-2015
7 JUN 1E-1937
12 TEAD3 1E-1250
13 TEAD1 1E-1185
14 TEAD4 1E-1140
16 TEAD 1E-1106
21 TEAD2 1E-907
53 TFAP2A 1E-171
57 TFAP2C 1E-162
77 GATA2 1E-122
79 GATA3 1E-121
1 TFAP2C 1E-74
2 TFAP2A 1E-62
3 TEAD1 1E-45
4 TEAD3 1E-39
5 TEAD 1E-35
6 TEAD4 1E-34
7 TEAD2 1E-28
9 TCF3 1E-14
12 LEF1 1E-10
J
Pluripotency
Mesoderm
Trophoblast
BMP4 d2
BMP4 d4
BMP4/IWP2 d2
BMP4/IWP2 d4
NANOG
EOMES
GATA3
TBXT
EGFR

**Supplementary Figure S1. A.** Principal component analysis (PCA) of RNA-seq data in hESCs differentiated into the trophoblast lineage with BMP4 in the presence and absence of IWP2. **B.** Venn diagram of up-regulated genes following treatment of hESCs with BMP4 with and without IWP2. **C.** Venn diagram of up-regulated genes following treatment of hESCs with BMP4 with and without IWP2 for 4 days. Table shows gene list enrichment analysis on the KEGG pathway database (2021) for the genes uniquely up-regulated with BMP4 alone. **D.** Epifluorescence images of TCF/LEF-eGFP reporter hESC line (HUES9) during trophoblast differentiation with BMP4, BMP4/IWP2, and BMP4/IWR1 to detect canonical WNT activation. **E.** Gene expression levels of mesoderm and WNT markers in TCF/LEF-eGFP reporter hESC line (HUES9) after trophoblast differentiation with BMP4, BMP4/IWP2, and BMP4/IWR1 by qPCR. Statistical analysis compared to BMP4 treatment. **F.** UMAP of annotated cells generated using the Early Embryogenesis Projection Tool (v2.1.2) (1). Samples were integrated and shown as red dots in the early embryogenesis projection tool (grey). **G.** UMAP with samples colored using the early embryogenesis projection tool's annotation prediction **(Table S2). H.** Pie chart showing the distribution of open chromatin sites throughout the annotated genomic locations in differentiated hESCs. **I.** Motif analyses using HOMER showing the significantly enriched DNA-binding motifs at accessible chromatin loci. **J.** Genome browser tracks showing chromatin accessibility peaks (ATAC-seq) for representative pluripotency, mesoderm, and trophoblast gene loci during treatment of hESCs with BMP4 and BMP4/IWP2. * $p \leq 0.05$, ** $p \leq 0.01$, *** $p \leq 0.001$, **** $p \leq 0.0001$.

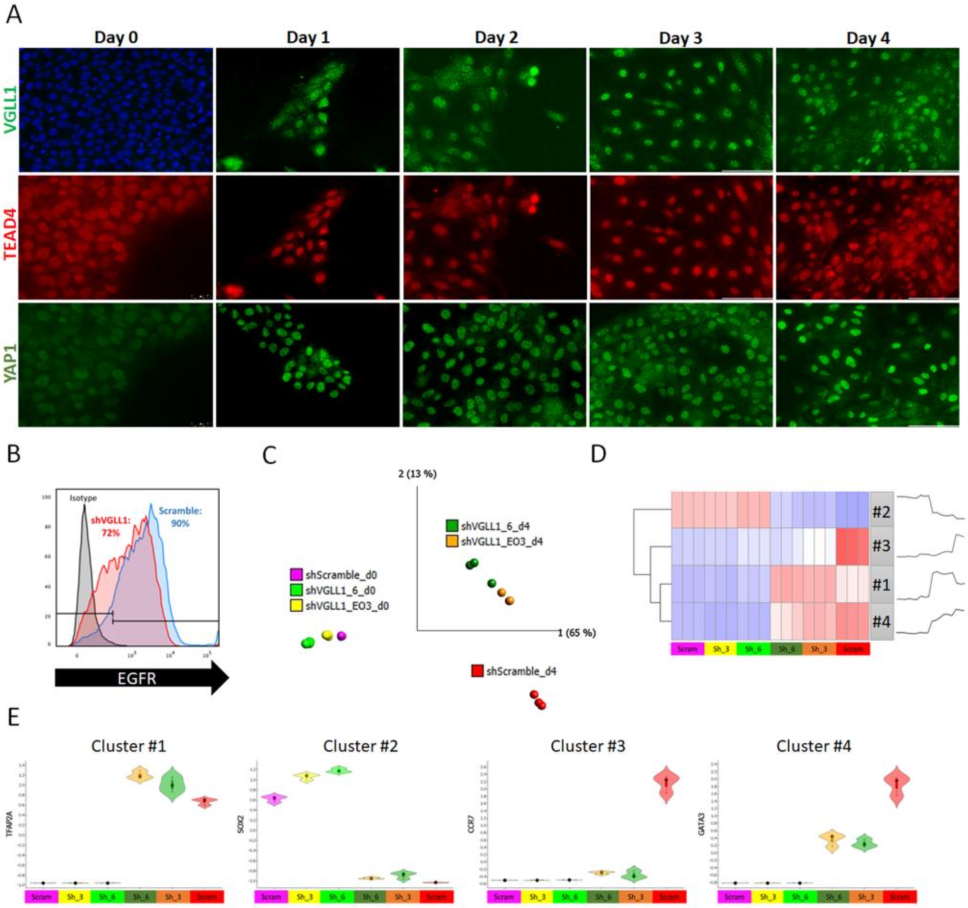


F

**Up-regulated genes delayed in shVGLL1**

| | Log(q-value) |
|---|---|
| Membrane Trafficking | -68.0 |
| protein catabolic process | -50.7 |
| membrane organization | -45.5 |
| intracellular protein transport | -42.6 |
| Signaling by Receptor Tyrosine Kinase | -37.9 |
| Signaling by Rho GTPases | -37.7 |
| Endocytosis | -34.8 |
| VEGFA VEGFR2 signaling | -34.8 |
| endosomal transport | -34.6 |
| regulation of cellular catabolic process | -34.3 |
| actin cytoskeleton organization | -34.3 |
| localization within membrane | -33.3 |
| endocytosis | -33.3 |
| response to hormone | -31.8 |
| Neutrophil degranulation | -31.0 |
| Lysosome | -30.3 |
| Growth factor signaling | -30.1 |
| Endoplasmic reticulum processing | -30.1 |
| cell migration | -28.7 |

**Down-regulated genes delayed in shVGLL1**

| | Log(q-value) |
|---|---|
| DNA metabolic process | -36.1 |
| Cell Cycle | -35.9 |
| chromatin remodeling | -35.8 |
| chromosome organization | -32.0 |
| regulation of DNA metabolic process | -22.1 |
| DNA replication | -22.0 |
| mitotic cell cycle | -22.0 |
| neuron projection morphogenesis | -21.2 |
| cell projection organization | -20.2 |
| modulation of chemical synaptic transmission | -19.1 |
| Endoderm differentiation | -17.6 |
| regulation of cell cycle process | -17.4 |
| brain development | -16.9 |
| behavior | -15.9 |
| negative regulation of cellular component organization | -15.3 |
| SUMO E3 ligases SUMOylate target proteins | -14.1 |
| embryo development | -13.6 |
| Neuronal System | -13.0 |
| protein localization to chromosome | -12.9 |
| phosphorylation | -12.6 |

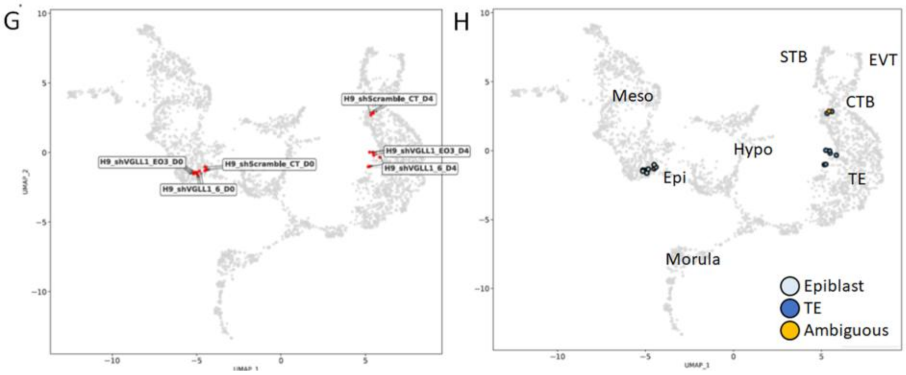

**Supplementary Figure S2. A.** Immunofluorescence staining for VGLL1, TEAD4, and YAP1 of hESCs during 4 days of BMP/IWP2 treatment. **B.** Cell surface expression of EGFR in Scramble and shVGLL1 hESCs at day 4 of BMP4/IWP2 treatment by flow cytometry. **C.** Principal component analysis (PCA) of RNA-seq data in Scramble and shVGLL1 hESCs. **D.** Affinity propagation (AP) analysis of the 7,284 DEGs in shVGLL1 hESC cells treated with BMP4/IWP2 compared to Scramble. **E.** Expression profiles of genes representative of each of the AP clusters. **F.** Gene ontology analysis of mis-regulated genes in shVGLL1 hESC cells treated with BMP4/IWP2 compared to Scramble. **G.** UMAP of annotated cells generated using the Early Embryogenesis Projection Tool (v2.1.2) (1). Samples were integrated and shown as red dots in the early embryogenesis projection tool (grey). **H.** UMAP with samples colored using the early embryogenesis projection tool's annotation prediction (**Table S2**).

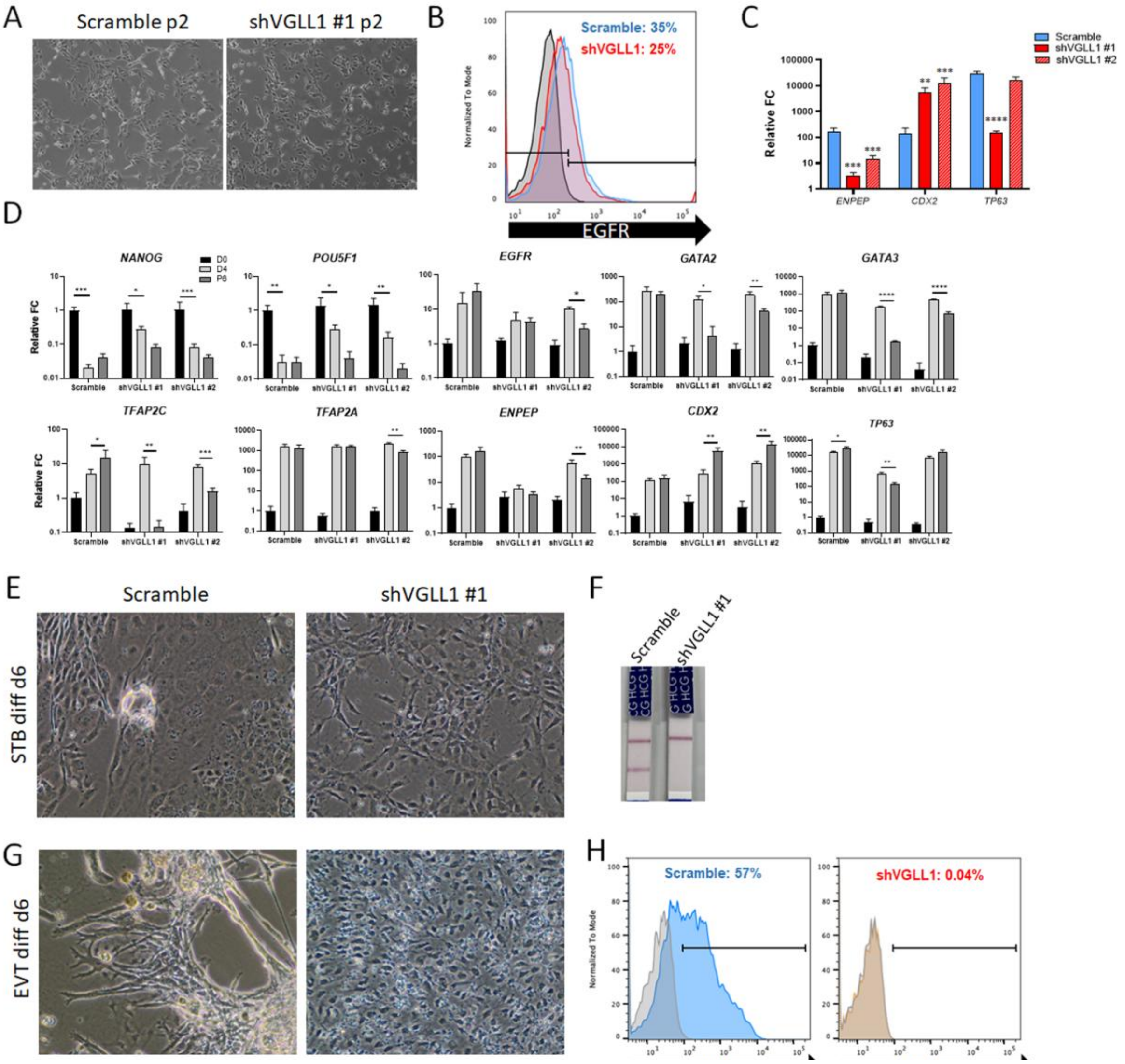


**Supplementary Figure S3. A.** Bright field images of Scramble and shVGLL1 hESCs at passage 2 of adaptation to hTSCs culture conditions. **B.** Cell surface expression of EGFR in Scramble and shVGLL1 hESCs at passage 2 of adaptation to hTSCs culture conditions by flow cytometry. **C.** Gene expression levels of early TE markers ENPEP and CDX2 and late trophoblast marker TP63 in Scramble and shVGLL1 hESC-derived TSCs by qPCR. **D.** Gene expression level comparison of various lineage markers between d0 hESCs, BMP4/IWP2-treated hESC (d4), and hESC-derived TSC (P6) in Scramble and shVGLL1 cells by qPCR. * $p \leq 0.05$, ** $p \leq 0.01$, *** $p \leq 0.001$, **** $p \leq 0.0001$. **E.** Bright field images of TSC-adapted Scramble and shVGLL1 cells differentiated into STB media for 6 days. **F.** hCG levels in spent media from TSC-adapted Scramble and shVGLL1 cells differentiated into STB media for 6 days. **G.** Bright field images of TSC-adapted Scramble and shVGLL1 cells differentiated into EVT media for 6 days. **H.** Cell surface expression of HLA-G in TSC-adapted Scramble and shVGLL1 cells differentiated into EVT media for 6 days by flow cytometry.

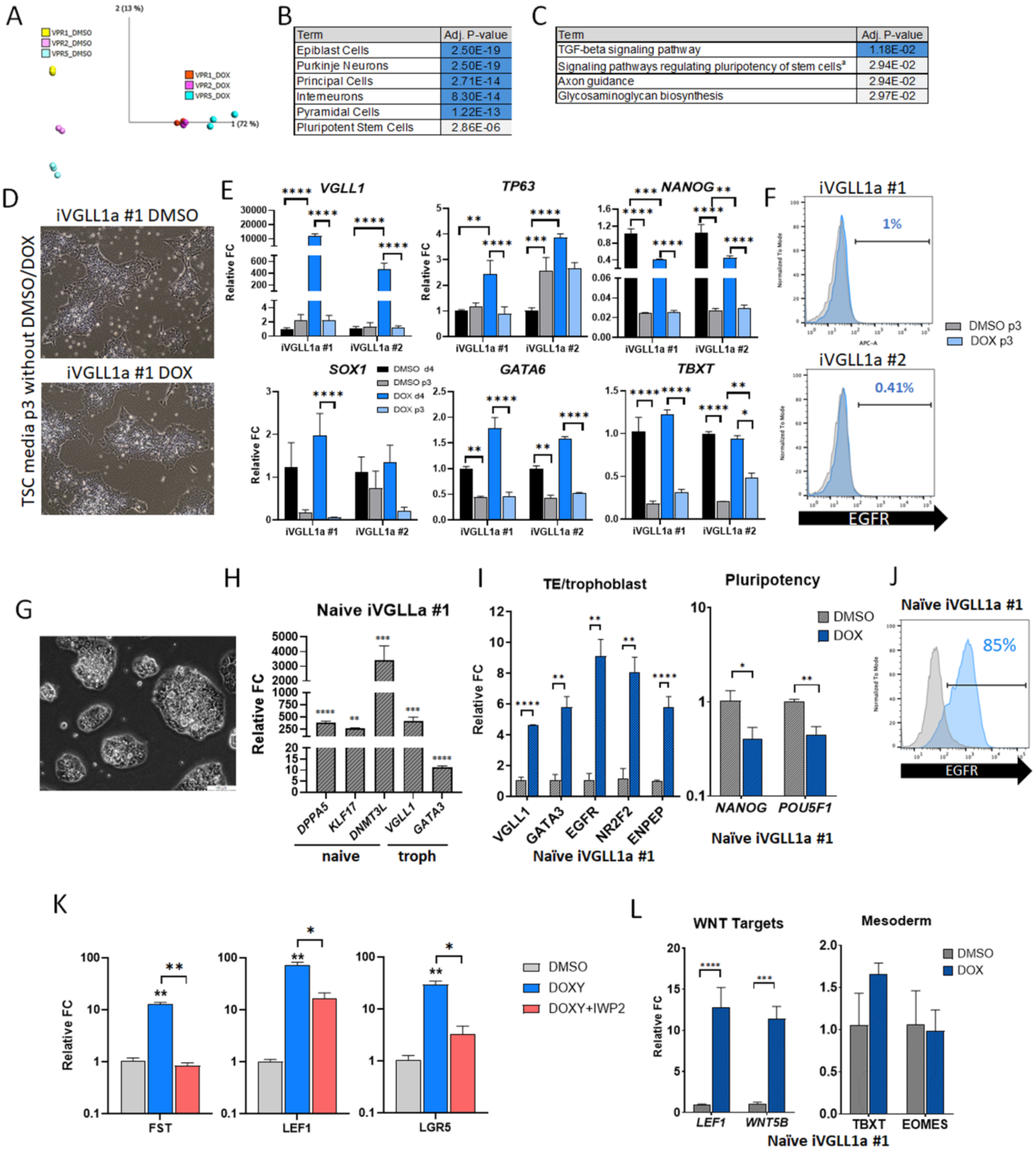


**Supplementary Figure S4. A.** Principal component analysis of RNA-seq data in DMSO- and DOX-treated iVGLL1a hESCs. **B.** Gene list enrichment analysis on the Panglao DB augmented database (2021) for 3,477 differentially down-regulated genes in DOX-treated iVGLL1a hESCs compared to DMSO. **C.** Gene list enrichment analysis on the KEGG pathway database (2021) for of 3,053 genes differentially up-regulated in DOX-treated iVGLL1a hESCs compared to DMSO, a=WNT, TGFb, MAPK, and PI3K-Akt signaling pathways. **D.** Bright field images of iVGLL1a clone #1 cells treated with either DMSO or DOX for 4 days in StemFlex media and subsequently cultured in TSC media for 3 passages without ectopic induction. **E.** Gene expression analysis of *VGLL1*, *TP63* (TSC marker), *NANOG* (pluripotency), *SOX1* (ectoderm), *GATA6* (endoderm), and *TBXT*

(mesoderm) in DMSO and DOX-treated cells after culture in TSC media for 3 passages without ectopic induction. **F.** Cell surface expression of EGFR in DMSO and DOX-treated cells after culture in TSC media for 3 passages without ectopic induction by flow cytometry. **G.** Bright field images of naive iVGLL1a hESCs adapted to PXGL media. **H**. Gene expression levels of naïve markers as well as TE/trophoblast markers, *VGLL1* and *GATA3* in naïve iVGLL1a hESCs compared to primed state cells by qPCR. **I.** Gene expression levels of TE/trophoblast and pluripotency markers in DOX-treated naïve iVGLL1a hESCs compared to DMSO by qPCR. **J.** Cell surface expression of EGFR in DMSO (grey) and DOX- treated (blue) naïve iVGLL1a hESCs by flow cytometry. **K.** Gene expression levels of canonical WNT target genes in DOX-treated iVGLL1a hESCs with and without IWP2 compared to DMSO by qPCR. **L.** Gene expression levels of WNT target genes, *LEF1* and *WNT5B*, and mesoderm markers, *EOMES* and *TBXT*, in naïve DOX-treated iVGLL1a hESCs compared to DMSO by qPCR. * $p \leq 0.05$, ** $p \leq 0.01$, *** $p \leq 0.001$, **** $p \leq 0.0001$.

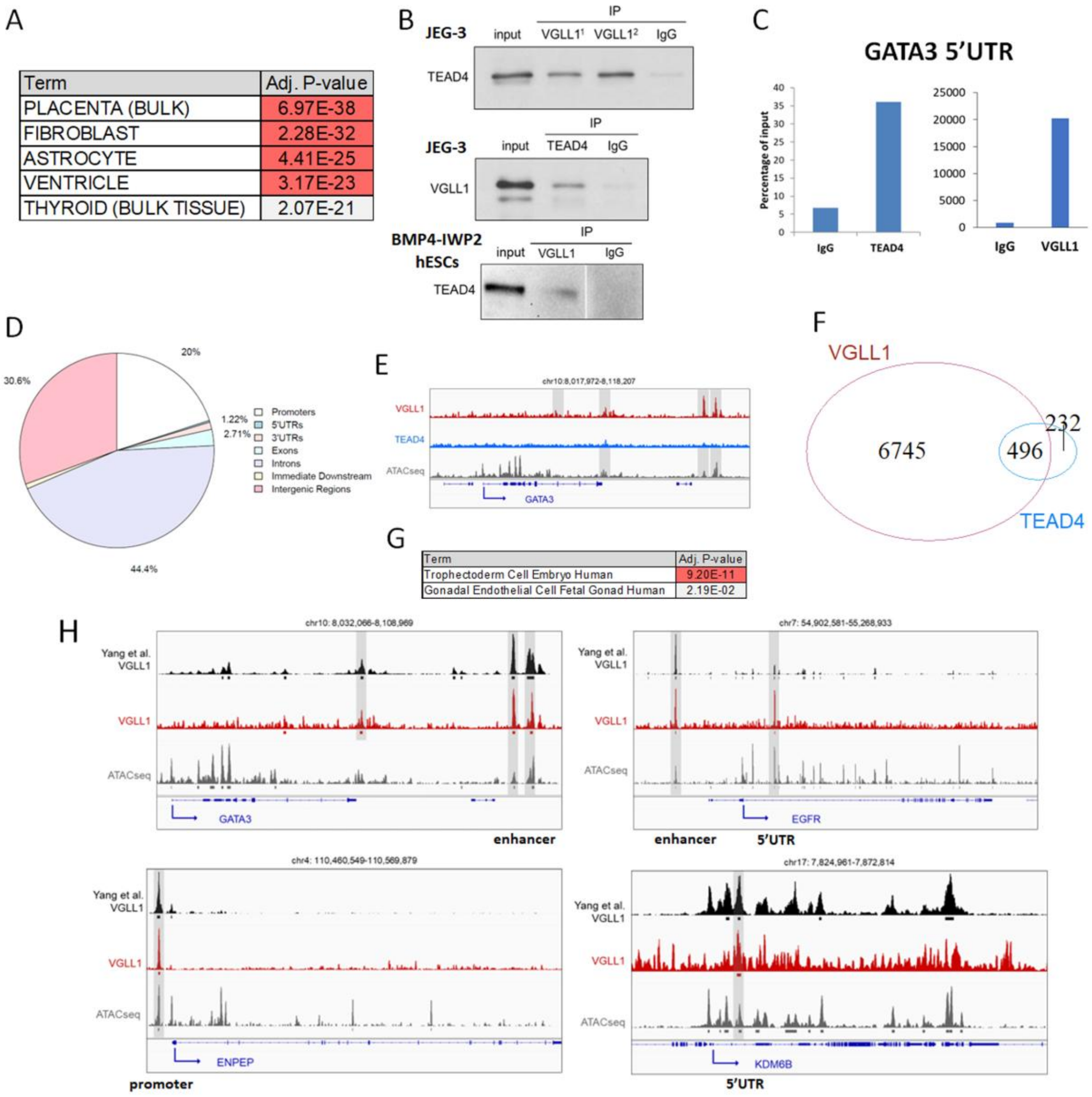


**Supplementary Figure S5. A.** Gene list enrichment analysis on the ARCHS4 Tissues database for 528 genes concordantly regulated by VGLL1 (down in shVGLL1 and up in DOX/IWP2-treated iVGLL1a hESCs). **B.** Western blot of TEAD4 and VGLL1 in JEG3 lysate and d4 BMP4-IWP2-treated hESCs co-immunoprecipitated with either VGLL1 or TEAD4 antibodies respectively. **C.** VGLL1 and TEAD4 ChIP-qPCR at a validated 5'UTR region of GATA3 in JEG3 cells. **D.** Pie chart showing the distribution of VGLL1-binding sites throughout the annotated genomic locations in differentiated hESC. **E.** Genome browser tracks showing VGLL1 and TEAD4 genomic enrichment peaks (ChIP-seq) and chromatin accessibility peaks (ATAC-seq) on GATA3 of BMP4/IWP2-treated hESCs at day 4 showing presence of a distal regulatory peak in VGLL1 but not TEAD4 tracks. **F.** Venn diagram of VGLL1 and TEAD4 uniquely annotated genomic enrichment peaks (ChIP-seq) in BMP4/IWP2-treated hESCs at day 4. **G**. Gene list enrichment analysis on the CellMarker (2024) database for 5,039 genes with VGLL1 occupancy (ChIPseq). **H**. Genome browser tracks showing shared VGLL1 genomic enrichment peaks between Yang et al. CUT&TAG (naïve TE cells) and our ChIP-seq data as well as chromatin accessibility peaks in our ATAC-seq (BMP4/IWP2-treated cells) on *GATA3, EGFR, ENPEP, KDM6B*.

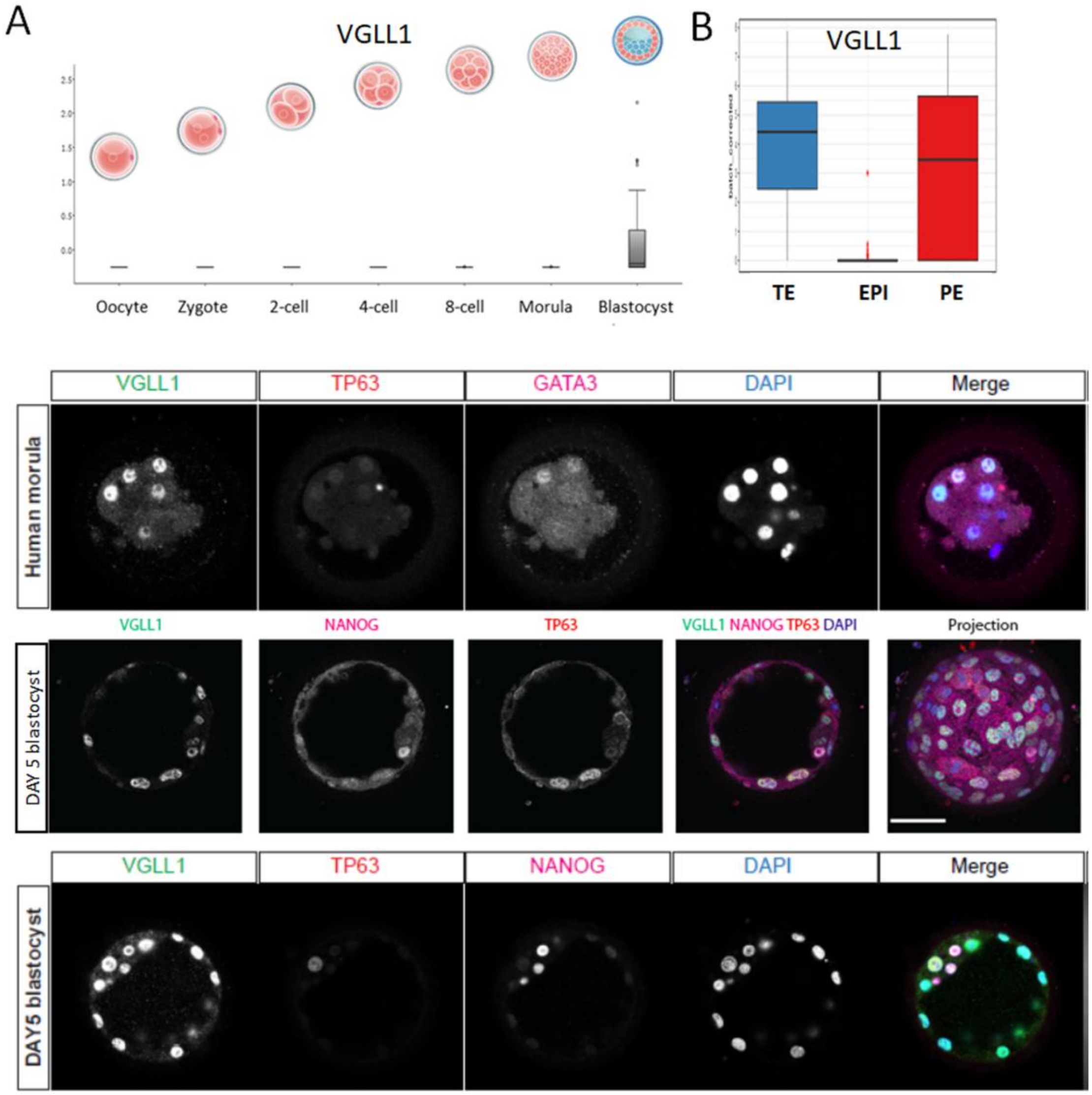


**Supplementary Figure S6. A.** Gene expression levels of VGLL1 during early embryo development (bulk RNA-seq data from Yan *et al.* 2013) (2). **B.** Gene expression levels of VGLL1 in pre-implantation blastocyst compartments (bulk RNA-seq data from Blakeley *et al.* 2015) (3). **C.** Immunofluorescence staining of a human morula with VGLL1, TP63, and GATA3 antibodies. **D**. Immunofluorescence staining of two day-5 blastocysts with VGLL1, TP63, and NANOG.

**Supplementary Methods**

**Culture of human embryonic stem cells.** Human embryonic stem cells H9/WA09 (H9) and HUES9TOPGFP were cultured and passaged in mTeSR Plus (StemCell Technologies, cat. no. 100-1130) on Geltrex (Gibco, cat. no. A14133-02)-coated plates as previously described (4). shRNA H9 hESC lines were previously derived and maintained in 10 μg/ml puromycin (Sigma Aldrich, cat. no. P8833-25MG) (5).

**Derivation of dCas9-VPR VGLL1 gRNAs (iVGLL1a) H9 hESCs.** Five RNA guides for VGLL1 activation were designed using the sgRNA Designer tool by the Broad Institute (6) and cloned into the Golden Gate pMA Cas9 Guide vector system (7), modified to incorporate an Eif1a-Bsd antibiotic resistance cassette for selectable incorporation into a mammalian cells. Each of the guides were cloned into the U6 promoter driven cloning site on the plasmid, and sequence verified. Plasmids containing DOX-inducible dCas9-VPR (gift from Dr. Church) (8), VGLL1-specific guide RNAs, and TOPGFP.mC (Addgene, cat. no. 35491) (9) were introduced in H9 ESCs using the Amaxa Nucleofector II electroporation system following manufacturer instructions (setting B-016). Briefly, 1 million cells were collected, resuspended and mixed with 2 μg of plasmid DNA before nucleofection using the Lonza Human Stem Cell Nucleofector Kit 1 (Lonza, cat. no. VPH-5012). Cells were immediately re-plated, and selection started 48 hours later. iVGLL1a cell lines were maintained with 2 μg/mL puromycin and 8 μg/mL blasticidin S (Gibco, cat. no. A1113903). Ectopic expression of VGLL1 was induced with 2 μg/mL doxycycline (Millipore Sigma, cat. no. D3447-500MG) for 4 days.

**Naïve conversion of primed human embryonic stem cells.** iVGLL1a H9 hESCs were converted to a naïve pluripotency state following previously established protocols (10–12). Briefly, primed hESCs were adapted to culture in 5% $O_2$ in the XVIVO Incubation System/Stem Cell Workstation (Biospherix) for 3 passages. Naïve conversion was induced using chemical reset with 1 μM PD0325901 (R&D Systems, cat. no. 4192), 10 ng/mL human recombinant LIF (PeproTech, cat. no. 300-05), and 2 μM valproic acid (R&D Systems, cat. no. 2815) in MEF-conditioned N2B27 media for 3 days followed by expansion with PXGL (1 μM PD0325901 (R&D Systems, cat. no. 4192), 2 μM XAV 939 (R&D Systems, cat. no. 3748), 2 μM Gö 6983 (R&D Systems, cat. no. 2285), 10 ng/mL human recombinant LIF (PeproTech, cat. no. 300-05)) in MEF-conditioned N2B27 media. Naïve hESCs were differentiated into TE cells following an established protocol (13). Briefly, 10ng/mL BMP4 (R&D Systems, cat. no. 314-BP-010), 2 μM A83-01 (SelleckChem, cat. no. S7692), and 2 μM PD0325901 (R&D Systems, cat. no. 4192) were added to N2B27 media for 1 day followed by 2 μM A83-01 (SelleckChem, cat. no. S7692), 2 μM PD0325901 (R&D Systems, cat. no. 4192), and 1 μg/mL Pyridone 6 (SelleckChem, cat. no. S6789), for 2 more days.

**Syncytiotrophoblast (STB) and extravillous trophoblast (EVT) differentiation of trophoblast stem cells (TSCs).** hPSC-derived TSCs were differentiated into STB ad EVT following previously published protocols (14, 15). STB differentiation was verified by morphology and hCG secretion in the media using HCG Urine DipStrips (CLIAwaived Inc., cat. no. CLIA-2471). EVT differentiation was verified by morphology and cell surface expression of HLA-G (EXBIO, cat. no. 1P-292-C100) by flow cytometry as previously described (16).

**Immunohistochemistry.** Formalin-fixed, paraffin-embedded placental tissue sections from three 5-week placentas were stained with anti-KDM6B (Abcam, cat. no. ab38113) and Goat anti-Rabbit IgG secondary antibody conjugated with AlexaFluor 488 (Invitrogen, cat. no. A11008) following previously published protocol (20). Fifteen embryos were thawed according to recommendations from Bourn Hall Clinic, the in vitro fertilization clinic coordinating donations using the Blast thaw kit (Origio, cat no. 10542010A), following the manufacturer's instructions and cultured in Gx-TL medium (Vitrolife, cat. no. 10172) for 4 hours prior to fixation in 4%

paraformaldehyde (Electron Microscopy Sciences, cat. no. 15710) in PBS, pre-chilled at 4°C. Differentiated hESCs were also fixed in 4% paraformaldehyde and stained as previously described (5). The following antibodies were used: rabbit anti-VGLL1 (Sigma Aldrich, cat. no. HPA042403-100UL), mouse anti-TEAD4 (Abcam, cat. no. ab58310), rabbit anti-YAP1 (Abcam, cat. no. ab52771), rabbit anti-KDM6B (Abcam, cat. no. ab38113), mouse anti-NANOG (Thermo Fisher, cat. no. MA1-017) or goat anti NANOG (R&D Systems, cat. no. AF1997), goat anti-GATA3 (R&D Systems, cat. no AF2605), mouse anti-p40 (ΔNp63-specific antibody, clone BC28; Biocare Medical, cat. no. ACI3066A), Alexa FluorTM 488 goat anti-rabbit IgG (Thermo Fisher, cat. no A-11008), Alexa FluorTM 594 goat anti-mouse IgG (Thermo Fisher, cat. no A-11005), Alexa Fluor™ 647 donkey anti-mouse IgG (Thermo Fisher, cat. no. A-31571), and Alexa Fluor™ 594 donkey anti-goat IgG (Thermo Fisher, cat. no. A-11058) Embryos were imaged on a Leica SP5 inverted confocal microscope (Leica Microsystems). Cells and placentas were imaged on an ECHO Rebel microscope (ECHO). Immunofluorescence images of KDM6B and VGLL1 in embryos were quantified using Fiji and Imaris software (v10.2.0) for 3D visualization of inside and outside cells, following a previously established protocol (17). Briefly, nuclei in immunofluorescently stained embryos were automatically segmented based on DAPI or Hoechst signals, and fluorescence intensities of KDM6B and VGLL1 were measured within the segmented nuclei.

**Flow cytometry.** Flow cytometry was conducted using live cells as previously described (4). Briefly, single cell suspensions were stained with EGFR-APC (Biolegend cat. no. 352906), or APC-conjugated mouse IgG (BioLegend, cat. no. 400120) antibodies, in flow cytometry buffer (1% FBS, 2% BSA, and 0.03% sodium azide) for 1 hour at 4°C. Cells were then washed three times in PBS and fixed with 2% PFA (Electron Microscopy Sciences, cat. no. 15710). Flow cytometry analysis was performed on a Fortessa X20 flow cytometer (BD Biosciences). Data were visualized using FlowJo Software (v10).

**Assay for Transposase-Accessible Chromatin (ATAC)-sequencing.** Nuclei extraction and ATAC-seq library prep ware performed according to previously published protocol (18). The bcbio-nextgen (v1.2.8) ATAC-seq pipeline was used to process the ATAC-seq reads. The nucleosome free fraction was normalized using deeptools (v 3.5.1) (19) bamCoverage by CPM. One sample with a lower signal quality than its replicates, as determined by limma-voom multi-dimensional scaling plots, was eliminated. Bigwigs for replicates were combined using wiggletools (v 1.2.11) (20) mean to obtain one average track from biological replicates. Genomic tracks were visualized using IGV Viewer (v2.11.7). Heatmaps were generated using deeptools computeMatrix and plotHeatmap. Binding profiles were generated deeptools computeMatrix and plotProfile. Consensus peaks were annotated using ChIPpeakAnno (v 3.32.0) (21) with EnsDb.Hsapiens.v86 (v 2.99.0). Chromosome regions were annotated using ChIPpeakAnno assignChromosomeRegion with TxDb.Hsapiens.UCSC.hg38.knownGene (v 3.16.0). Differential expression analysis was carried out using edgeR (v 3.40.2) (22) with limma-voom normalization (23). HOMER (v 5.1) (24) findMotifsGenome.pl was then used to identify transcription factor binding motifs. Mesoderm and TE genes for signal intensity analysis are listed in **Suppl. Table S7.**

**RNA isolation, cDNA preparation, and quantitative real-time PCR.** Total RNA was purified using NucleoSpin isolation kit (Macherey-Nagel, cat. no. 740955.250) and RNA concentration measured using NanoDrop 2000C spectrophotometer (ThermoScientific). cDNA was prepared from total RNA using the PrimeScript RT-Kit (Takara Bio, cat. no. RR037A). Quantitative real-time PCR (qPCR) was performed using TB Green Premix Ex Taq (Takara bio, cat. no. RR420A) on a QuantStudio5 thermocycler (AppliedBiosystems). The primer sequences used are listed in **Suppl. Table S8**. Relative expression of each transcript was calculated using ΔΔCT method, normalized to 18S rRNA (25). Statistical analysis and graphs were generated using GraphPad Prism 10. Bar chart data display mean fold change and standard deviation. Student's t tests were performed to determine significance between two groups. Two-way ANOVA was performed when analyzing significant differences between three or more groups. All qRT-PCR experiments were done using three replicates and statistical tests were performed using log transformed fold change. Level of significance is represented with asterisks.

**Co-immunoprecipitation and western blot.** Co-immunoprecipitation was performed on nuclear fractions obtained using the NE-PER Nuclear extraction kit (Thermo Scientific, cat. no. 78833) following the manufacturer's protocol on 20µL pellets of ~2 million cells. After harvesting, 200µL of ice-cold CER I (cat. no. 78833A) and 2µL of protease inhibitor (Halt, cat. no. 1861281) was added to the cell pellet, vortexed, and incubated for 10 min. on ice. 11µL of CER II (cat. no. 78833B) was then added followed by a brief vortex and a 1-minute incubation. Lysate was centrifuged for 5 min. at 16,000xg at 4°C and supernatant removed (cytosolic fraction). The remaining nuclear pellet was then resuspended in 100µL of NER (cat. no. 78833C) and 1µL of protease inhibitor. After 40 minutes of incubation, with 15-second vortexing at 10-minute intervals, the nuclear fraction was then centrifuged at 16,000xg for 10 min. at 4°C. 95µl of supernatant was collected and stored at -80°C. 105µL of IP lysis buffer (Pierce, cat. no. 87787) was added to bring the total volume to 200µL. Pre-clearing was done with 20µL of agarose beads (Cell Signaling Technology, cat. no. 9863S) for 45 min. at 4°C on a rocker. After centrifuging at 14,000xg at 4°C for 10 minutes, supernatant was transferred into a new microcentrifuge tube and aliquoted as follows: 10µL for input, 95µL for targeted precipitation, and 95µL for IgG control. Input was stored at -80°C. Cleared lysates were incubated in 2µg of anti-VGLL1 antibody (Sigma Aldrich, cat. no. HPA042403), anti-TEAD4 antibody (Abcam, cat. no. ab58310), or anti-IgG antibody (Sigma Aldrich, cat. no. NI01) overnight rotating at 4°C. Agarose beads (20 µL) were added and incubated at 4°C for 80 min. Samples were centrifuged at 14,000xg for 2 min. at 4°C and washed three times with lysis buffer. Immunoprecipitated proteins were eluted using 20µL of 2x Laemmli Sample Buffer (BIO-RAD, cat. no. 1610737) + 5µL of 2-Mercaptoethanol (BIO-RAD, cat. no. 1610710). Input and both bound and unbound fractions were denatured at 95°C for 10 min. Samples were loaded into separate wells of a Mini-PROTEAN TGX Gel (BIO-RAD, cat. no. 4561093) and ran at 120V for 80 min. in 1x Running Buffer. Transfer was done on using a Trans-Blot Turbo RTA PVDF Transfer Kit (BIO-RAD, cat. no. 1704272) in 1x Transfer Buffer. After transfer, the membrane was blocked in 2% milk in TBS-T for 60 min. at room temperature rocking followed by three washes for 10 min. in TBS-T. Membrane was incubated overnight at 4°C rocking with primary mouse anti-TEAD4 antibody (Abcam, cat. no. ab58310) for VGLL1-co-IP and rabbit anti-VGLL1 antibody (Sigma Aldrich, cat. no. HPA042403) for TEAD4 co-IP, diluted 1:1000 in 2% milk/TBS-T. After three washes, membrane was incubated on a rocker at room temperature for 60 min. with secondary goat anti-mouse HRP antibody (Thermo Scientific, cat. no. 31430) or goat anti-rabbit HRP (Thermo Scientific, cat. no. 31460) diluted 1:1000 in 2% milk/TBS-T. The membrane was imaged using a ChemiDoc MP Imaging System (BIO-RAD, cat. no. 170-8280) with SuperSignal West Dura Extended Duration Substrate (Thermo Scientific, cat. no. 34075). Images were analyzed using Image Lab (BIO-RAD).

**RNA sequencing and analysis.** Total RNA extraction with mirVana (Ambion, cat. no. AM1561) and library preparation for RNA-sequencing were performed as previously described (26). Quality control was performed using FastQC (v.0.12.1) and multiQC (v.1.19). Two samples with a lower signal quality than their replicates were eliminated. Reads were mapped to GRCh38.p14 (GENCODE release 45) using STAR (v.2.7.9a) (27) and annotated using featureCounts (subread v.2.0.6). Ensembl genes without at least three samples with 10 or more reads were removed from analysis. Normalization and differential expression analysis was performed using the R (v.4.4.0) package DESeq2 (v.1.46.0) (28). Genes with an adjusted p-value < 0.05 and log2 fold change > 1 were considered differentially expressed. In iVGLL1a models, adjusted p-value < 0.05 and log2 fold change > 0.5 were used to include subtle but biologically relevant changes. Heatmaps, PCA, and violin plots were generated with Qlucore. Statistical analysis was performed using a two-group comparison or multi-group comparison with q-value <0.05. Gene list enrichment analysis was performed with Enrichr on multiple database as specified in the text (29).

**Chromatin Immunoprecipitation (ChIP)-sequencing.** Chromatin immunoprecipitation was performed following a previously published protocol (30). Briefly, cells were crosslinked with 1% formaldehyde (Pierce, cat. no. 28906) for 10 minutes at room temperature on a horizontal rocker. Crosslinking was stopped with 2.5 M glycine. Cells were collected into 15mL conical tubes and centrifuged at 1200rpm for 5 minutes at 4°C. Cell pellets were washed in PBS two times and cell pellets were stored in -80°C until ChIP. Frozen cell pellets were resuspended in cell lysis buffer with protease inhibitors and incubated at 4°C for 10 minutes. Following centrifugation, nuclear pellets were resuspended in nuclei lysis buffer with protease inhibitors and incubated at 4°C for 10 minutes. Covaris shearing buffer was added and chromatin was sheared in microtube-500 AFA (Covaris, cat. no. 520185) using Covaris E220 Focused-Ultrasonicator with settings 5% duty, PiP 140, 15 minutes in 30 second bursts, cycles 200. Sodium chloride concentration was adjusted to 150mM and Triton-X to 1% following shearing. Pre-clearing was done with agarose beads (Cell Signaling Technology, cat. no. 9863S) for 30 minutes at 4°C on a rocker. Input was collected and stored in -20°C. Cleared lysates were incubated with 6 µg of mouse anti-TEAD4 (Abcam, cat. no. ab58310) or rabbit anti-VGLL1 antibody (Sigma Aldrich, cat. no. HPA042403-100UL) overnight at 4°C on a rocker. Agarose beads (50 µL) were added and incubated at 4°C for 1 hour on a rocker. Lysates were centrifuged at 14000rpm for 5 minutes at 4°C and washed five times. DNA-protein-antibody complexes were eluted with 0.1 M sodium bicarbonate and 1% SDS, then vortexed briefly and centrifuged at 7500rpm for 2 minutes at room temperature. Reverse crosslinking for eluted and input DNA was done with 3 µg/mL RNAse A (ThermoScientific, cat. no. EN0531) and 0.3 M sodium chloride (Invitrogen, cat. no. AM9759) on a heat block at 67°C for 5 hours. Proteinase K (Invitrogen, cat. no. 100005393) was added (0.2 mg/mL) and incubated at 45°C overnight. DNA purification was performed with Monarch PCR and DNA Cleanup kit (NEB, cat. no. T1030) and submitted for library preparation and sequencing conducted at the IGM Genomics Center, University of California, San Diego, La Jolla, CA.

**ChIP-seq analysis.** Adapters were trimmed from reads using Trim Galore (v0.6.10) -q 20 --fastqc --paired. Trimmed reads were aligned to GRCh38 using bowtie2 (v2.4.1) with setting -p 24. samtools (v1.10) were used to sort and index bam files. Peaks were called using MACS2 (v2.2.6) callpeak with settings -f BAMPE -g 2.7e+9 -q 0.05 --keep-dup all --call-summits. DiffBind (v3.16.0) was used to find consensus peaks among replicates using minOverlapmembers = 2. BigWig files were generated with deeptools (v3.5.5) bamCoverage with settings --binSize 20 --smoothLength 60 --effectiveGenomeSize 2913022398 –normalizeUsing CPM --extendReads 150 --centerReads --scaleFactorsMethod None --minMappingQuality 30 --ignoreForNormalization chrM, then replicates were averaged using bigwigAverage with settings -bs 20. Heatmaps were generated with deeptools computeMatrix reference-point --referencePoint center -b 1000 -a 1000 -missingDataAsZero -refPointLabel 0 and visualized using plotHeatmap --kmeans 2. ChIPpeakAnno (v3.14.2) was used to annotate peaksets. Motif analysis was performed using HOMER (v5.1) findMotifsGenome.pl with -size 200. Pie charts of genomic annotations were generated with ChIPpeakAnno pie1 function and genomic element distribution gED. Genomic tracks were visualized using IGV Viewer (v2.11.7). Computationally intensive sequencing analyses were done on the San Diego Supercomputer Center (2025): Expanse through the University of California San Diego (31).

**CUT&RUN-qPCR.** CUT&RUN assay (Cell Signaling Technology, cat. no. 86652) was performed following manufacturer's protocol on 150,000 cells per replicate. Briefly, cells were bound to Concanavalin A Beads, permeabilized with 0.05% digitonin, and incubated on a nutator overnight with 1 µg of anti-H3K27me3 antibody

(Diagenode, cat. no. C15410195). Bound DNA was digested with pAG-MNase activated with calcium chloride and released with a 10 minute incubation without shaking at 37°C. DNA was purified with Monarch PCR & DNA cleanup kit (cat. no. T1030). Primers for qPCR on CUT&RUN samples are listed in **Suppl. Table S8**. Negative control primers targeting the ACTB promoter were purchased from ActiveMotif (cat. no. 71023). CUT&RUNqPCR data was analyzed by 2-△△CT method (32). Fold enrichment was calculated relative to undifferentiated cells and statistical significance was assessed by two-way ANOVA in GraphPad Prism 10 (v10.6.0).

**Supplementary Table S1. Differentially expressed genes of H9 hESCs treated with BMP4 or BMP4-IWP2 by bulk RNA-seq.** Separate tabs contain DEGs for the following comparisons: BMP4 at d2 vs d0, BMP4 at d4 vs d0, BMP4-IWP2 at d2 vs d0, and BMP4-IWP2 at d4 vs d0.

**Supplementary Table S2. Cell annotation based on the human embryo reference tool.** Separate tabs contain annotation for 1. hESCs do0, BMP4 d2, BMP4-IWP2 d2, BMP4 d4, BMP4-IWP2 d4; 2. Scramble and shVGLL1 clones d0 and BMP4-IWP2 d4.

**Supplementary Table S3. Differential peak analysis of H9 hESCs treated with BMP4 or BMP4-IWP2 by ATAC-seq.** Separate tabs contain differential peaks for the following comparisons: BMP4 at d2 vs d0, BMP4 at d4 vs d0, BMP4-IWP2 at d2 vs d0, and BMP4-IWP2 at d4 vs d0. The last two tabs list differential peaks between BMP4 and BMP4-IWP2 at d2 and d4, respectively.

**Supplementary Table S4. Differentially expressed genes between Scramble and shVGLL1 hESCs at d4 of BMP4-IWP2 treatment by bulk RNA-seq.**

**Supplementary Table S5. Differentially expressed genes in iVGLL1a hESCs treated with Dox or DOX-IWP2 compared to DMSO control by bulk RNA-seq.** The last tab list genes concordantly regulated by VGLL1: failed to be up-regulated in shVGLL1 hESCs and up-regulated in DOX-IWP2-treated iVGLL1a hESCs.

**Supplementary Table S6. Differential peak analysis of H9 hESCs treated with BMP4-IWP2 for 4 days by ChIP-seq for TEAD4 and VGLL1.** Tabs list unique and common peaks in VGLL1 and TEAD4 ChIP-seq.

**Supplementary Table S7. TE and mesoderm genes**

| **Mesoderm*** | **TE**** |
|---|---|
| MIXL1 | EGFR |
| TBXT | TFAP2A |
| VIM | TFAP2C |
| LEFTY1 | GATA2 |
| SOX17 | GATA3 |
| GATA6 | CDX2 |
| MESP1 | CCR7 |
| EOMES | NR2F2 |
| TBX6 | ENPEP |
| LHX1 | TACSTD2 |
| FOXA2 | DAB2 |
| FGF18 | KRT1 |
| PDGFRA | ABCG2 |

*gene list compiled from references (33–35)

** gene list compiled from references (13, 36)

**Supplementary Table S8. Primer list**

**qRT-PCR**

| **Primers** | **Forward** | **Reverse** |
|---|---|---|
| 18S | CGCCGCTAGAGGTGAAATTCT | CGAACCTCCGACTTTCGTTCT |
| ASCL2 | CACTGCTGGCAAACGGAGAC | AAAACTCCAGATAGTGGGGGC |
| CDX2 | TTCACTACAGTCGCTACATCACC | TTGATTTTCCTCTCCTTTGCTC |
| DNMT3L | TTCTGGATGTTCGTGGACAA | ACATCTGGGATGGTGACTGG |
| DPPA5 | TGAAAGATCCAGAGGTGTTC | ACTGGTTCACTTCATCCAAG |
| EGFR | CTAAGATCCCGTCCATCGCC | GGAGCCCAGCACTTTGATCT |
| ELF5 | AGTCTGCACTGACATTTTCTCATC | CAGAAGTCCTAGGGGCAGTC |
| ENPEP | TCCCAAAGAATACGGAGCACT | TCATGGGGACAGACTTCTCAA |
| EOMES | TGCCTACCAAAACACCGATATTAC | GGAGAATCCGTGGGAGATGG |
| FST | GCTGCTCTGCCAGTTCATGG | CGTTCTTCGCTTGACGGAGC |
| GATA2 | CCTGTAGTTCCTGCCCCTCT | AGCTGCCGACTCCCAGA |
| GATA3 | AGGGACGTCCTGTGCGAACT | GGTCTGGATGCCTTCCTTCTTCA |
| GATA6 | TCTACAGCAAGATGAACGGCC | AGGTGGAAGTTGGAGTCATGG |
| GCM1 | TTTTTCCAGTCAAAGGGAGAGCA | AGGAAGGTGCTGTGTTCACTT |
| KLF17 | CTGCCTGAGCGTGGTATGAG | TCATCCGGGAAGGAGTGAGA |
| LEF1 | TTATCCAGGCTGGTCTGCAA | AGGCAGCTGTCATTCTTGGA |
| LGR5 | GGATGTTGCTCAGGGTGGACT | ATTCGGGAGCAGCTGACTGA |
| NANOG | CCTATGCCTGTGATTTGTGG | CTTTGGGACTGGTGGAAGAA |
| NR2F2 | GCCATAGTCCTGTTCACCTCA | AATCTCGTCGGCTGGTTG |
| POU5F1 | TGGGCTCGAGAAGGATGTG | GCATAGTCGCTGCTTGATCG |
| SOX1 | CTCACTTTCCTCCGCGTTGCT | TGCCCTGGTCTTTGTCCTTCA |
| TBXT | AGTTGAGGACAGCAGGTTTTA | GCTGATTGTCTTTGGCTACT |
| TFAP2A | GTTACCCTGCTCACATCACTAG | TCTTGTCACTTGCTCATTGGG |
| TFAP2C | GAAGAGGACTGCGAGGATCG | GCTGATATTCGGCGACTCCA |
| TP63 | CTGGAAAACAATGCCCAGA | AGAGAGCATCGAAGGTGGAG |
| VGLL1 | CTCCCGGCTCAGTTCACTATAA | CCCAGTGGTTTGGTGGTGTA |
| WNT5B | CCAACTCCTGGTGGTCATTAGC | TGGGCACCGATGATAAACATC |

**ChIP/CUT&RUN-qPCR**

| **Primers** | **Forward** | **Reverse** |
|---|---|---|
| EGFR 5'UTR | TAAGTGCCCAGAAGAGCAACA | GCAGGTGAGGTTGGAATGCT |
| GATA3 5'UTR | GGCGCCGTCTTGATACTTT | TCTCTCTCTCTCCCTCTCTCAG |
| GATA3 dist enh | GAATCCTGTCCTCCACAGCG | GGTTGGGGGAGAGAACAAGC |
